# Sympathetic Stimulation Can Compensate for Hypocalcaemia-Induced Bradycardia in Human and Rabbit Sinoatrial Node Cells

**DOI:** 10.1101/2024.08.30.610432

**Authors:** Moritz Linder, Tomas Stary, Gergő Bitay, Norbert Nagy, Axel Loewe

## Abstract

Regular activation of the heart originates from cyclic spontaneous depolarisations of sinoatrial node cells (SANC). Variations in electrolyte levels, commonly observed in haemodialysis (HD) patients, and the autonomic nervous system (ANS) profoundly affect the SANC function. Thus, we investigated the effects of hypocalcaemia and sympathetic stimulation on SANC beating rate (BR).

The β-adrenergic (β-AR) signalling cascade, as described by Behar et al., was incorporated into the SANC models of Severi et al. (rabbit) and Fabbri et al. (human). Simulations were conducted across various extracellular calcium ([Ca^2+^]_o_) (0.6 to 1.8 mM) and isoprenaline concentrations [ISO] (0 to 1000 nM) for a sufficient time to allow transient oscillations to equilibrate and reach a limit cycle. The β-AR cell response of the extended models was validated against new Langendorff-perfused rabbit heart experiments and literature data.

The extended models revealed that decreased [Ca^2+^]_o_ necessitated an exponential-like increase in [ISO] to restore the basal BR. Specifically, at 1.2 mM [Ca^2+^]_o_, the Severi and Fabbri model required 15.5 and 7.3 nM [ISO], respectively, to restore the initial BR. Further reduction of [Ca^2+^]_o_ to 0.6 mM required 60.0 and 41.7 nM [ISO] to compensate for hypocalcaemia. A sudden loss of sympathetic tone at low [Ca^2+^]_o_ resulted in extreme bradycardia or even loss of automaticity within seconds.

These findings suggest that hypocalcaemic bradycardia can be compensated for by an elevated sympathetic tone. The integration of the β-AR pathways led to a logarithmic BR increase and offers insights into potential pathomechanisms underlying sudden cardiac death (SCD) in HD patients.

**Key Points:**

- We extended the sinoatrial node cell (SANC) models of Severi et al. (rabbit) and Fabbri et al. (human) with the β-adrenergic (β-AR) signalling cascade Behar et al. described.
- Simulations were conducted across various extracellular calcium ([Ca^2+^]_o_) (0.6 to 1.8 mM) and isoprenaline concentrations [ISO] (0 to 1000 nM) to mimic conditions in haemodialysis patients.
- An exponential-like increase in [ISO] compensated for hypocalcaemia-induced bradycardia in both models, while inter-species differences lead to more sensitivity of the extended Fabbri model towards hypocalcaemia and increased sympathetic tone.
- The extended models may help to further understand the pathomechanisms of several cardiovascular diseases affecting pacemaking, such as the high occurrence of sudden cardiac death (SCD) in chronic kidney disease (CKD) patients.

## Introduction

The natural cardiac pacemaking function is initiated by periodic excitations of sinoatrial node cells, orchestrated by the intricate interplay of the calcium (Ca^2+^) and membrane currents, known as the coupled clock mechanism. This involves precise regulation of the sensitive balance of inward and outward transmembrane currents during the diastolic depolarisation (DD) phase, as well as complex interaction of both effects throughout the action potential (AP). Various factors, including the extracellular milieu and the autonomic nervous system (ANS), regulate sinus node electrophysiology, thereby influencing the beating rate (BR) [1]. By precisely regulating electrolyte concentrations in the blood and extracellular milieu, the kidneys are vital for maintaining homeostasis. However, in individuals suffering from chronic kidney disease (CKD), the renal system fails to keep the electrolyte levels within these narrow ranges, contributing to a 14-fold increase of the risk of sudden cardiac death (SCD) compared to patients with cardiovascular disease and normal kidney function [2]. Long term studies in haemodialysis (HD) patients with implantable loop recorders revealed that all observed SCDs occurred after progressive bradycardia followed by asystole rather than the traditional SCD mechanisms ventricular tachycardia or ventricular fibrillation [3]. Thus, attention turned to bradycardia and asystole as important pathomechanisms, which might be explained by the fluctuations in electrolyte concentrations, caused by HD or renal failure.

Loewe et al. investigated the effect of altered electrolyte levels on the sinus node pacemaking, highlighting that especially hypocalcaemia slows down the BR [4]. Furthermore, in patients with end-stage renal disease elevated plasma catecholamines, enhanced sensitivity to norepinephrine and increased muscular sympathetic nerve activity were found [5], [6]. CKD was also associated with a reduced heart rate response to sympathetic stimulation and it was found that an abrupt reduction of sympathetic tone acutely precedes SCD in a rat model [7].

In this study, we test the hypothesis that an increased sympathetic tone can compensate for hypocalcaemia-induced bradycardia to a certain extent, whereas a sudden loss under hypocalcaemic conditions could lead to severe bradycardia and SCD in CKD patients. We studied this in computational models of rabbit and human sinoatrial node cells (SANC). Current computational models barely allow for simulation of graded ANS effects, such as β-adrenergic (β-AR) stimulation on membrane and sarcoplasmic reticulum (SR) currents. There is an attempt at linear interpolation of conductance increases and gate shifts by Stary et al. [8] and a phenomenological approach by Maltsev et al. [9]. Demir et al. proposed a cascade in which only cyclic adenosine monophosphate (cAMP) modulated target ion channels [10], and Himeno et al. developed a guinea pig SANC model without consideration of subsarcolemmal space, resulting in low intracellular Ca^2+^ levels during local Ca^2+^ release [11]. Thus, we extended the SANC models of Severi et al. (rabbit) [12] and Fabbri et al. (human) [13] with the β-AR signalling cascade, which also takes the adenylyl cyclase (AC)-cAMP-protein kinase A (PKA) interplay into account, as suggested by Behar et al. [14]. The β-AR cell response of the extended model was validated by experiments with Langendorff-perfused rabbit hearts under the influence of various isoprenaline concentrations [ISO]. This allowed us to investigate the compensatory effect of [ISO] on hypocalcaemia-induced bradycardia and inter-species differences in mammalian pacemaking [15].

## Methods

We analysed the concentration-dependent compensatory effect of various [ISO] in reference to decreasing extracellular calcium concentrations ([Ca^2+^]_o_) in the *in silico* SANC models of Severi et al. (rabbit) [12] and Fabbri et al. (human) [13]. The model definitions were acquired from the CellML [16] and the openCARP [17] repository. The original model formulations allow simulating SANC electrophysiology either in basal conditions or under the influence of 1000 nM [ISO]. To consider graded changes of the sympathetic tone, modelling of the β-AR signalling cascade, based on the work of Behar et al. [14], was adjusted and integrated into these models.

Figure 1 shows a schematic illustration of the AC-cAMP-PKA signalling cascade after sympathetic stimulation. In general, isoprenaline (ISO) binding to the β-AR receptor or calmodulin (less pronounced) activates AC, which transforms adenosine triphosphate (ATP) into the second messenger molecule cAMP. To model the effect of ISO after stimulation of the β-AR receptor, ATP and cAMP levels were introduced in the model. The increase in cAMP directly affects the hyperpolarisation activated “funny” current (I_f_) by binding to the ion channel and activates the catalytic subunit of PKA. This leads to phosphorylation of the L-type Ca^2+^ current (I_CaL_) channel, sodium/potassium pump (I_NaK_), slow delayed rectifier K^+^ current (I_Ks_) channel, the ryanodine receptor (RyR) and phospholamban (PLB) regulating the sarcoplasmic/endoplasmic reticulum Ca^2+^-adenosine triphosphatase (SERCA). The phosphorylation was modelled by the introduction of a PKA activity and PLB level, which increased channel conductance and shifted activation curves.

**Figure 1:**
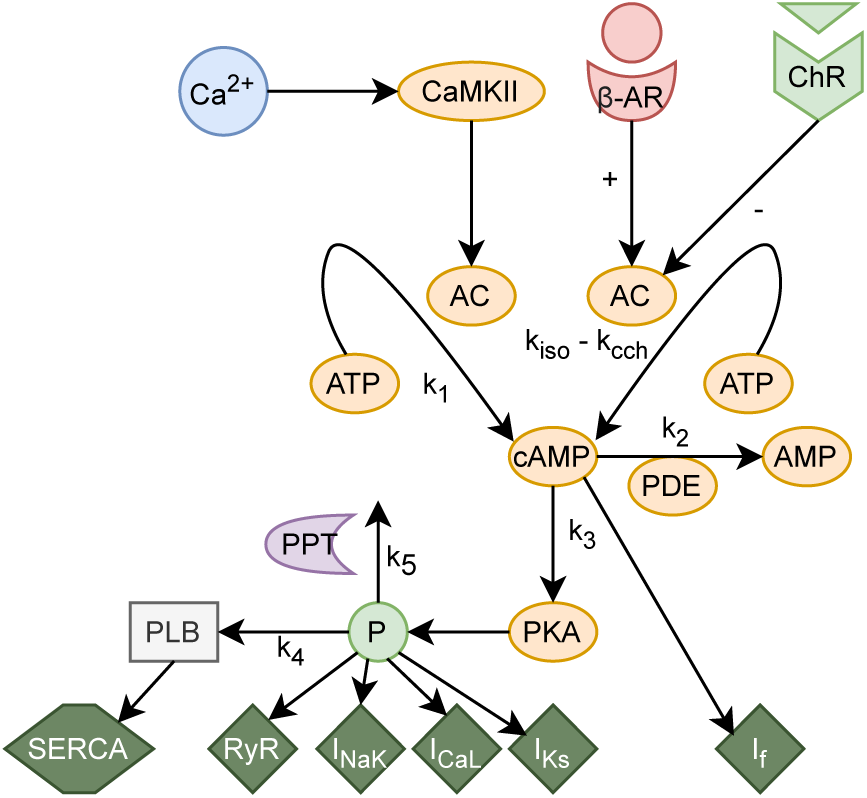
Schematic illustration of the β-AR signalling cascade. Adenylyl cyclase (AC) is activated byβ-adrenergic receptor (β-AR) and Ca2+/calmodulin-dependent protein kinase II (CaMKII) and deactivated by cholinergic receptor (ChR) stimulation. Activated AC transforms adenosine triphosphate (ATP) into second messenger molecule cyclic adenosine monophosphate (cAMP), which itself activates the catalytic subunit of protein kinase A (PKA). cAMP directly inffuences the “funny” current channel (If), while PKA phosphorylates several targets, including phospholamban (PLB), whose phosphorylation level regulates SERCA. Furthermore, two restraining mechanisms regulate the phosphorylation and cAMP level. Protein phosphatase (PPT) removes phosphate groups from proteins, whereas phosphodiesterase (PDE) breaks phosphodiester bonds of cAMP.

The Behar-Yaniv (BY) model is based on the Maltsev-Lakatta (ML) rabbit SANC model [18] and was extended by the addition of AC-cAMP-PKA signalling under β-AR stimulation by Yaniv et al. [19]. Caused by the different emphasis of the role of Ca^2+^ and membrane clocks, which were heavily discussed in the past, both the ML and BY model are based on a pronounced Ca^2+^ cycling. In contrast, the Severi model considers I_f_ as an additional potent driver of pacemaker function. Therefore, the initial parameters and current formulations of the BY and Severi rabbit SANC model differ, which required changes in the signalling cascade prior to the integration into the Severi model detailed below. Due to a scarcity of experimental data for human SANC, the β-AR signalling cascade was first adjusted and implemented into the Severi (rabbit) SANC model and later upscaled and integrated into the Fabbri (human) SANC model. The changes detailed below included the introduction of the adjusted cAMP-PKA relationship (Eq. (1) as well as ATP and PLB levels.

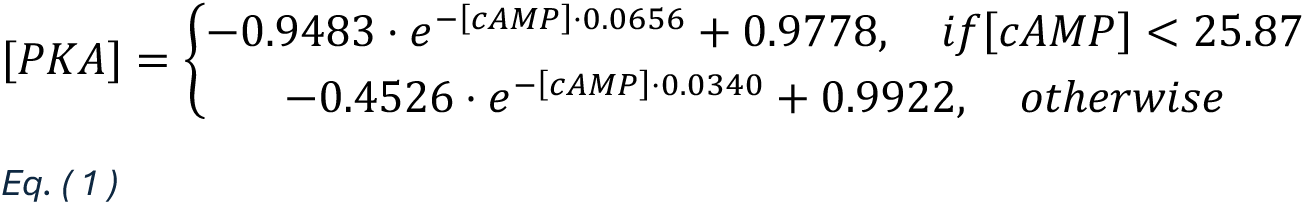

More specifically, the modulation of the activation gate of I_f_ followed a Hill-like equation and was adopted from the BY model [19]. To ensure a similar behaviour between the extended Severi and BY model for the basal concentration of cAMP, the Hill coefficient and a multiplicative factor (Fabbri model) of the equation were optimised using the minimize function with sequential least squares programming of the *SciPy* library in python [20]. The modifications are listed in Table 1.

**Table 1:**
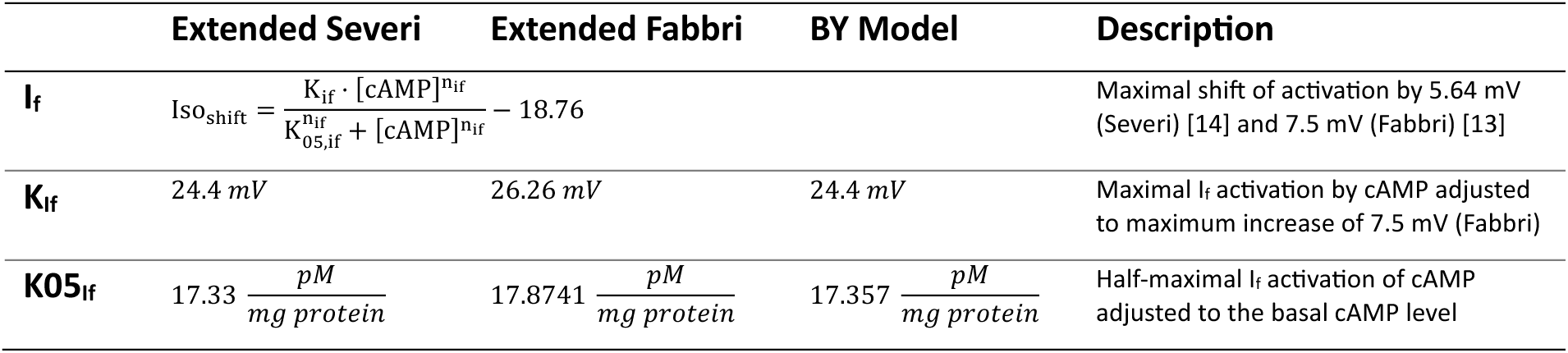
β-AR effects on If in the different SANC models.

The level of PKA-dependent phosphorylation modulates I_CaL_ [21]. Thus, in the BY model, a maximal I_CaL_ conductance increase of 80 % was implemented [14]. Based on the responses of the original Severi and Fabbri model to 1000 nM [ISO], the modulation of I_CaL_ in the extended models was split into an increase in conductance of 23 %, a shift of the activation curve by -8 mV and an inverse slope factor of the activation variable dL_∞_ decrease by 31 % (only Severi model). The increase in conductance was adjusted by altering the Hill coefficient and multiplicative factor of the equation, similar to the optimisation of I_f_ described above. The shift of the activation gate was assumed to respond to PKA like the I_f_ gating variable to cAMP and the inverse slope factor was assumed to follow a linear decrease. The modifications are listed in Table 2.

**Table 2:**
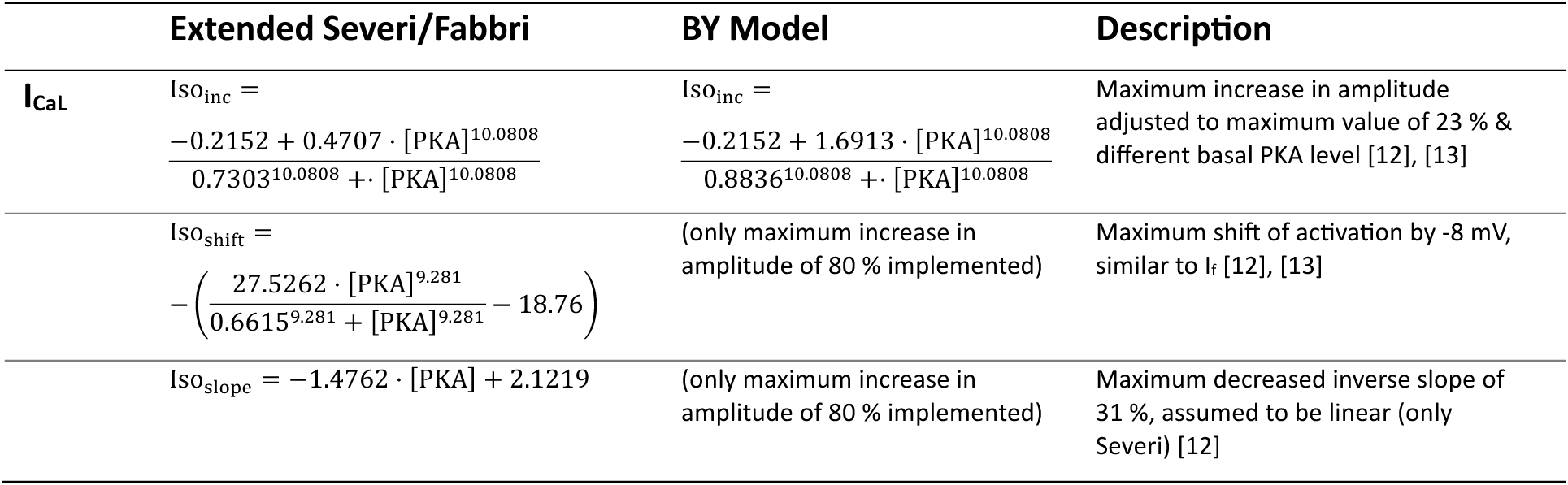
β-AR effects on ICaL in the different SANC models.

In the BY model, the PKA-mediated phosphorylation of I_Ks_ and I_NaK_ was not implemented. To balance large increases in the overshoot (OS) potential and action potential duration (APD) for high [ISO], the modulation of these two currents was necessary in the extended Severi and Fabbri model. At 1000 nM [ISO], the conductance of I_Ks_ and I_NaK_ were increased by 20 % while only the activation curve of I_Ks_ was shifted by -14 mV. These modulations were assumed to follow a similar Hill curve as I_CaL_. All changes introduced to I_Ks_ and I_NaK_ are listed in Table 3.

**Table 3:**
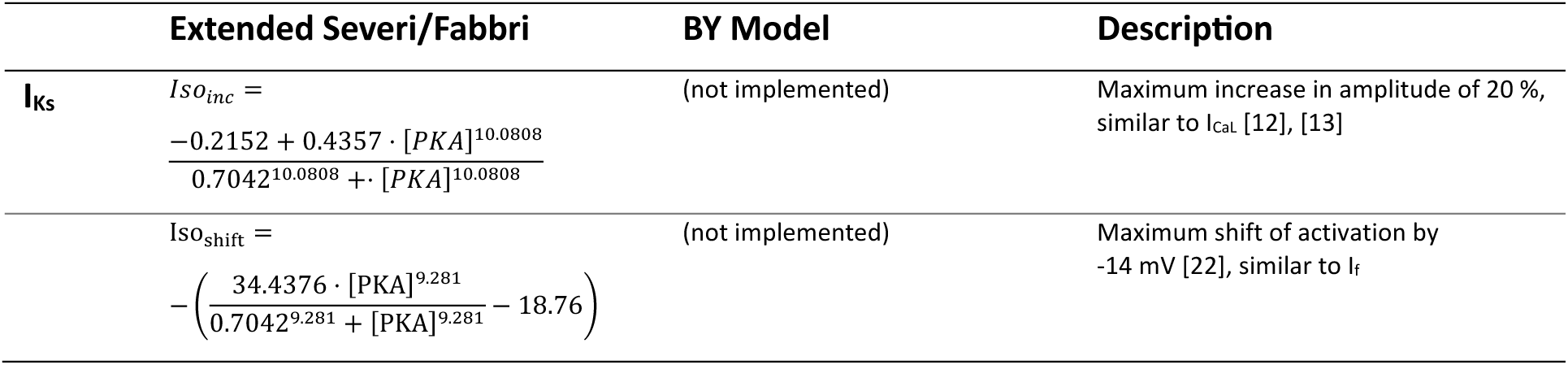
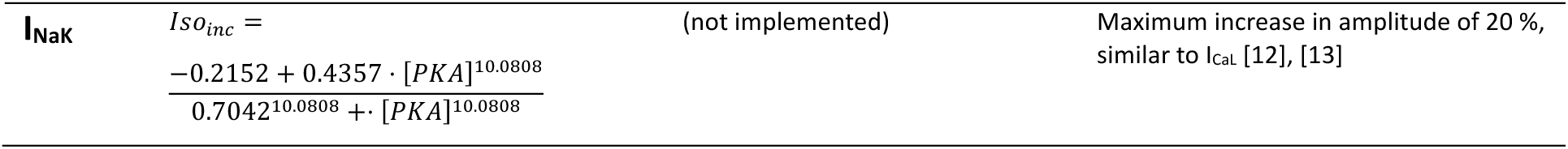
β*-AR effects on IKs and INaK in the different SANC models*.

The influence of the phosphorylation on the Ca^2+^ release by RyR with respect to PKA was modified to compensate for the slightly different basal level of PKA. Table 4 lists the introduced changes of the Ca^2+^ release by RyR. The effects on SERCA were implemented phenomenologically and adopted from [14].

**Table 4:** β-AR effects on RyR Ca^2+^ release in the different SANC models.

|  | Extended Severi/Fabbri | BY Model | Description |
| --- | --- | --- | --- |
| **koCa** | $koCa = koCa_{max} \cdot \left( RyR_{min} - \frac{RyR_{max} \cdot [PKA]^{n_{RyR}}}{K_{05,RyR}^{n_{RyR}} + [PKA]^{n_{RyR}}} - 1 \right)$ | | RyR phosphorylation [19] |
| **K05<sub>RyR</sub>** | 0.6829 | 0.7 | Half-maximal RyR activation of PKA, adjusted to different basal PKA level |

The degradation of PLB phosphorylation was adjusted in the extended Fabbri model to ensure a PLB activity level between 0 and 1. Table 5 lists the introduced changes in the PLB degradation. The phosphorylation of PLB regulating SERCA was implemented based on the experimental data from [23].

**Table 5:** β-AR effects on PLB degradation in the different SANC models.

|  | Extended Severi | Extended Fabbri | BY Model | Description |
| --- | --- | --- | --- | --- |
| | $k5 = \frac{k_{PP1} \cdot PP1 \cdot PLB_p}{k_{PP1,PLB} + PLB_p}$ | | | Degradation of PLB phosphorylation by protein phosphatase (PPT) |
| **k<sub>PP1</sub>** | $23,575 \frac{1}{\mu M \cdot min}$ | $23,850 \frac{1}{\mu M \cdot min}$ | $23,575 \frac{1}{\mu M \cdot min}$ | Maximal PPT activity |
| **K<sub>PP1,PLB</sub>** | 0.06967 | 0.07457 | 0.06967 | Half-maximal PPT activity |

Finally, different Ca^2+^-related association and dissociation constants (kb_CM_ & kf_CM_) in the BY and the extended models, along with differences in the fractional occupancy of calmodulin by Ca^2+^ inside the myoplasm (fCMi) required adaptations in the dynamic calculation of the cAMP concentration. Therefore, the constant defining the maximal AC activation by intracellular Ca^2+^ concentration ([Ca^2+^]_i_) (K_Ca_) was altered in both models and a multiplicative factor K_fab_=4.64 was introduced in the Fabbri model (Table 6). All model equations, constants and initial values are documented in the supplementary material.

**Table 6:**
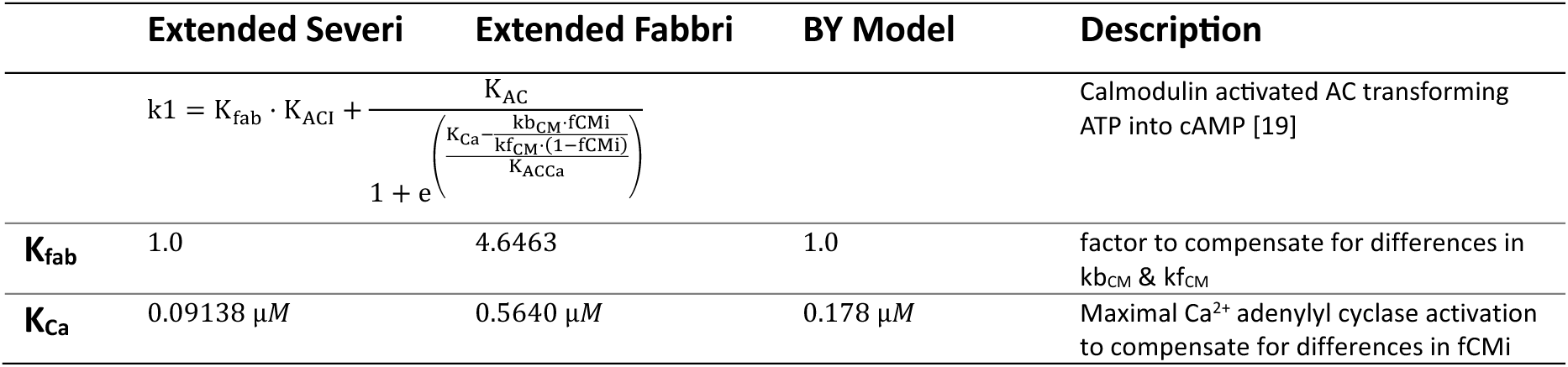
β-AR effects on cAMP generation in the different SANC models.

### Simulation Study

The models were solved using openCARP [17] with Rush-Larsen and forward Euler integration schemes for 1000 s, with the last 100 s simulated time considered during analysis. This allowed transient oscillations to equilibrate, resulting in a stable limit cycle. [Ca^2+^]_o_ was varied between 0.6 and 1.8 mM with a step size of 0.2 mM, whereas [ISO] ranged from 0 to 10 nM in steps of 2.5 nM, from 10 to 100 nM in steps of 25 nM and 100 to 1000 nM in steps of 250 nM.

To analyse the dominant pacemaking mechanisms that mediate the autonomic regulation via cAMP-PKA signalling, all simulations were performed again with specific ISO targets modelled as insensitive to phosphorylation, i.e., remaining in basal conditions. To assess the relative contribution of the various drivers to BR changes in relation to [Ca^2+^]_o_ and [ISO], the main currents were analysed after temporal integration during the AP phases, i.e. considering the charge transferred by them during a certain period. Lastly, for each AP cycle, the following markers were calculated: (a) maximum diastolic potential (MDP), defined as the most negative potential during repolarisation; (b) overshoot (OS), defined as the maximum potential during the AP cycle; (c) 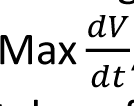, defined as the maximum of the first order derivative of the transmembrane voltage; (d) take off potential (TOP), defined as the transmembrane voltage measured at the time the voltage derivative 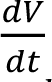 exceeds 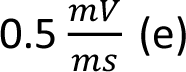 (e) action potential duration (APD) at 20, 50, 90 and 100 %, defined as time of 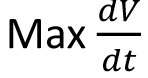 to the point were the AP amplitude repolarised to 20, 50, 90 or 100 %; (f) diastolic depolarisation rate 100 (DDR100), defined as the voltage change in the first 100 ms after MDP; (g) diastolic depolarisation duration (DD), defined as the time between MDP and TOP; (h) DD early, defined as the first 100 ms after MDP; (i) DD late, defined as DD without considering the first 100 ms after MDP.

### Langendorff-perfused Heart Experiments

All experiments were conducted in compliance with the Guide for the Care and Use of Laboratory Animals (USA NIH publication No 85-23, revised 1996) and conformed to Directive 2010/63/EU of the European Parliament. The protocols were approved by the Review Board of the Department of Animal Health and Food Control of the Ministry of Agriculture and Rural Development, Hungary (XIII./1211/2012). Animal studies were carried out in compliance of ARRIVE guidelines. Animals were obtained from a licenced supplier (Sano-Pikk Ltd, Hungary, Licence No.:PE/EA/00607/-2/2021).

New-Zealand white rabbits from both sexes were sacrificed by concussion after 400 IU heparin were injected intravenously. The excised hearts were mounted by the aorta on a Langendorff-apparatus and retrograde perfused with modified Krebs-Henseleit solution (KHB) at a constant pressure (80 Hgmm). The KHB solution contained (in mM): NaHCO_3_ 25; KCl 4.3; NaCl 118.5; MgSO_4_ 1.2; KH_2_PO_4_ 1.2; glucose 10; CaCl_2_ 1.8, having a pH of 7.4 ± 0.05 when gassed with 95% O_2_ + 5% CO_2_. The electrical activity as electrocardiogram (ECG) was obtained by using three lead custom-made electrodes and signal amplifier (Experimetria Ltd., Budapest, Hungary). Signal processing and analysis was carried out using HaemoSys (Experimetria Ltd., Budapest, Hungary). After control recordings various concentrations of ISO (Merck Life Science Ltd, Darmstadt, Germany) were employed, and the ECG R-R intervals were analysed. All experiments were carried out at 37 °C.

## Results

### Effects of Increased Sympathetic Tone

The extended Severi model in the basal state yielded a BR of 174.7 bpm (CL 343.4 ms) with APD90 and DD shares of 39.6 % (135.7 ms) and 48.5 % (166.3 ms), respectively. Based on current integration I_f_ and the sodium/calcium exchanger current (I_NaCa_) transferred the most charge during the complete DD. When increasing [ISO] to 10 nM, the BR was elevated by 11.7 %, while further increases to 100 nM resulted in an elevation of 23.6 %. At around 500 nM, a saturation effect was observed with a BR increase of 35.7 %, whereas the maximum increase was 36.0 % (174.9 to 237.9 bpm) at 1000 nM [ISO]. Selected currents of one complete cycle under basal conditions and the influence of 10 nM [ISO] are shown in Figure 2a,b.

**Figure 2:**
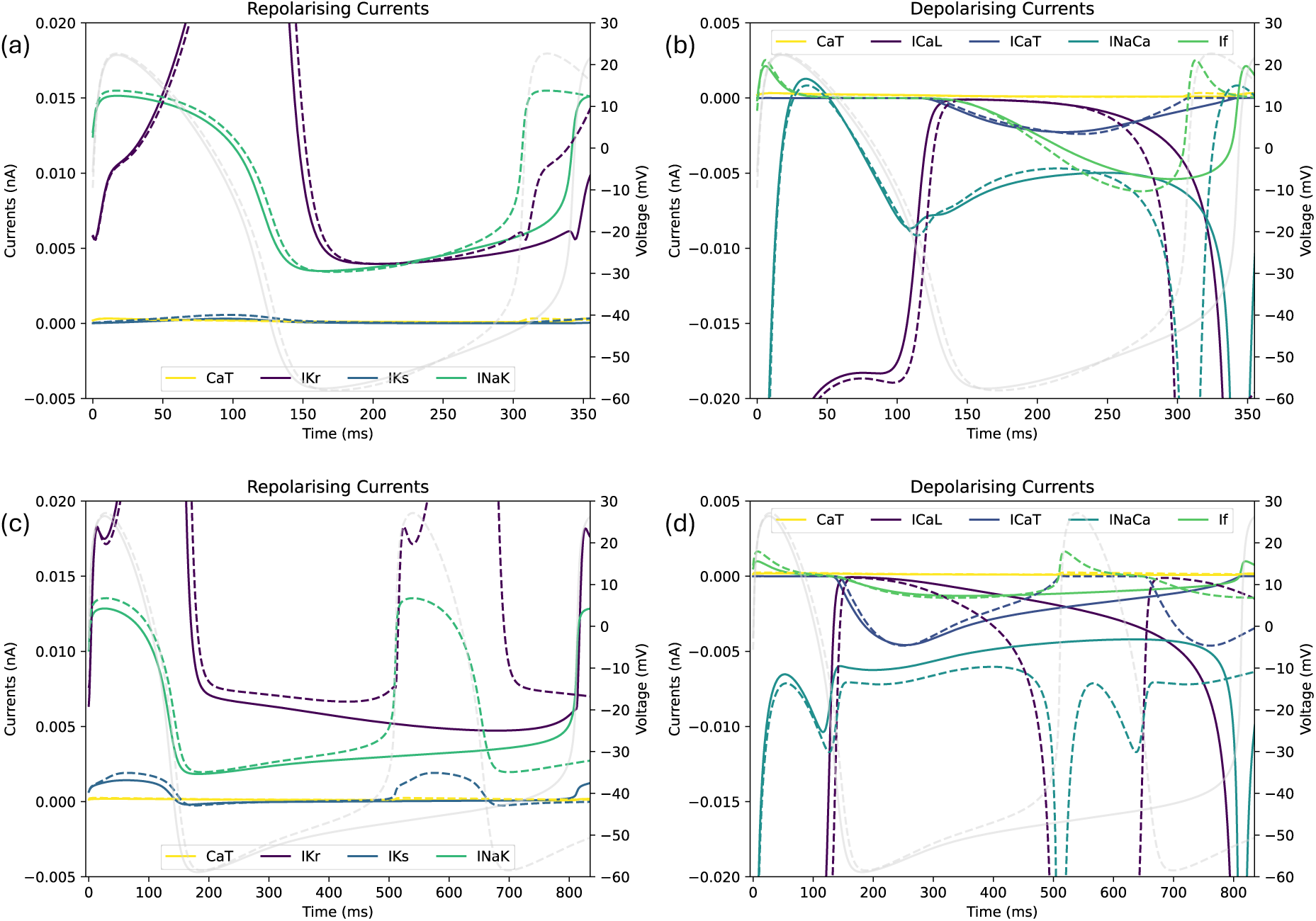
(a) selected repolarising and (b) depolarising currents of the extended Severi model during one AP cycle under basal conditions (solid) and after β-AR stimulation with 10 nM [ISO] (dashed). (c) selected repolarising and (d) depolarising currents of the extended Fabbri model during one AP cycle under basal conditions (solid) and after β-AR stimulation with 10 nM [ISO] (dashed). Transmembrane voltage in light grey (background). Note the different scales for the currents.

Simulations with the extended Fabbri model highlighted inter-species differences. In the basal state, the BR was 73.8 bpm (CL 813.0 ms) with APD90 and DD shares of 18.5 % (150.8 ms) and 76.2 % (619.9 ms), respectively. The influence of I_f_ was attenuated, which rebalanced the pacemaking control towards I_CaL_ and T-type Ca^2+^ channel current (I_CaT_). The influence of I_NaCa_ remained similar. The transferred charge during the first 100 ms after MDP was similar, with 1.1 and 1.2 pC for the extended Severi and Fabbri model, respectively. The attenuation of I_f_ during early DD was mainly compensated for by the increased contribution of I_CaT_. However, during late DD, the compensation for the reduced contribution of I_f_ switched to I_CaL_ with the transferred charge by I_CaT_ already declining due to starting inactivation of the channel.

Since the activation of I_CaL_ was controlled by voltage gates and a Ca^2+^ activated gate, I_CaL_ largely increased shortly after the elevated cytoplasmic Ca^2+^ concentration caused by the sarcoplasmic Ca^2+^ release of RyR. The delayed Ca^2+^ release extenuated the slope of I_CaL_ and delayed the abrupt downstroke of a main contributor to membrane depolarisation and AP initiation. Since repolarising currents (e.g., I_Kr_, I_Ks_ and I_NaK_) were similarly active in both models, the rebalanced contribution, from I_f_ to I_CaT_ and I_CaL_, led to a prolonged DD by 453.6 ms. This meant a higher total amount of charge transferred during DD, compared to the extended Severi model. Consequently, in the extended Fabbri model, the influence of the Ca^2+^ cycling was increased, the CL was increased, and therefore the BR was reduced. In contrast to DD, APD90 changed only slightly by a total of 15.1 ms.

Increasing [ISO] to 10 nM elevated the BR by 59.1 %, which already exceeded the maximum of the extended Severi model. Further increases of [ISO] to 100 nM resulted in elevations of 156.3 %. Whereas the saturation effect could be observed at around 250 nM [ISO], with an acceleration of 182.0 %, the maximum increase at 1000 nM was 183.3 % (73.8 to 209.1 bpm). Figure 2c,d shows selected currents under basal conditions and the influence of 10 nM [ISO] for the extended Fabbri model. Caused by elevated cAMP concentration and PKA activity level, the BR accelerated under sympathetic stimulation. In case of the extended Severi model, the cAMP level was elevated by 8.8 %, whereas PKA activity increased by 3.9 %. Similar increases were observed for the extended Fabbri model with elevations of 10.9 and 4.8 %, respectively.

To assess the ion charge transferred between the cell compartments during the complete AP cycle, selected currents were integrated and the relative contribution to depolarisation and repolarisation of these currents was calculated. For DD I_CaL_, I_CaT_, I_NaCa_ and I_f_ were considered. The raised cAMP concentration shifted the activation curve of I_f_ to more positive transmembrane voltages (Hill-equation with a maximum of 5.6 and 7.5 mV for 1000 nM [ISO] in the extended Severi and Fabbri model), which opened the voltage gate of the channel earlier during DD and increased the charge transferred during early DD. In the extended Severi model, the charge transferred by I_f_ during early DD was 0.3 pC (27.8 % of the charge of the selected currents) in basal conditions and increased to a contribution of 34.2 % under the influence of 10 nM [ISO]. For the extended Fabbri model, the basal contribution was lower with 0.1 pC (8.4 %), with a negligible increase of 0.2 % for 10 nM [ISO]. Figure 3 shows the relative contribution to re- and depolarisation of selected currents for several [ISO].

**Figure 3:**
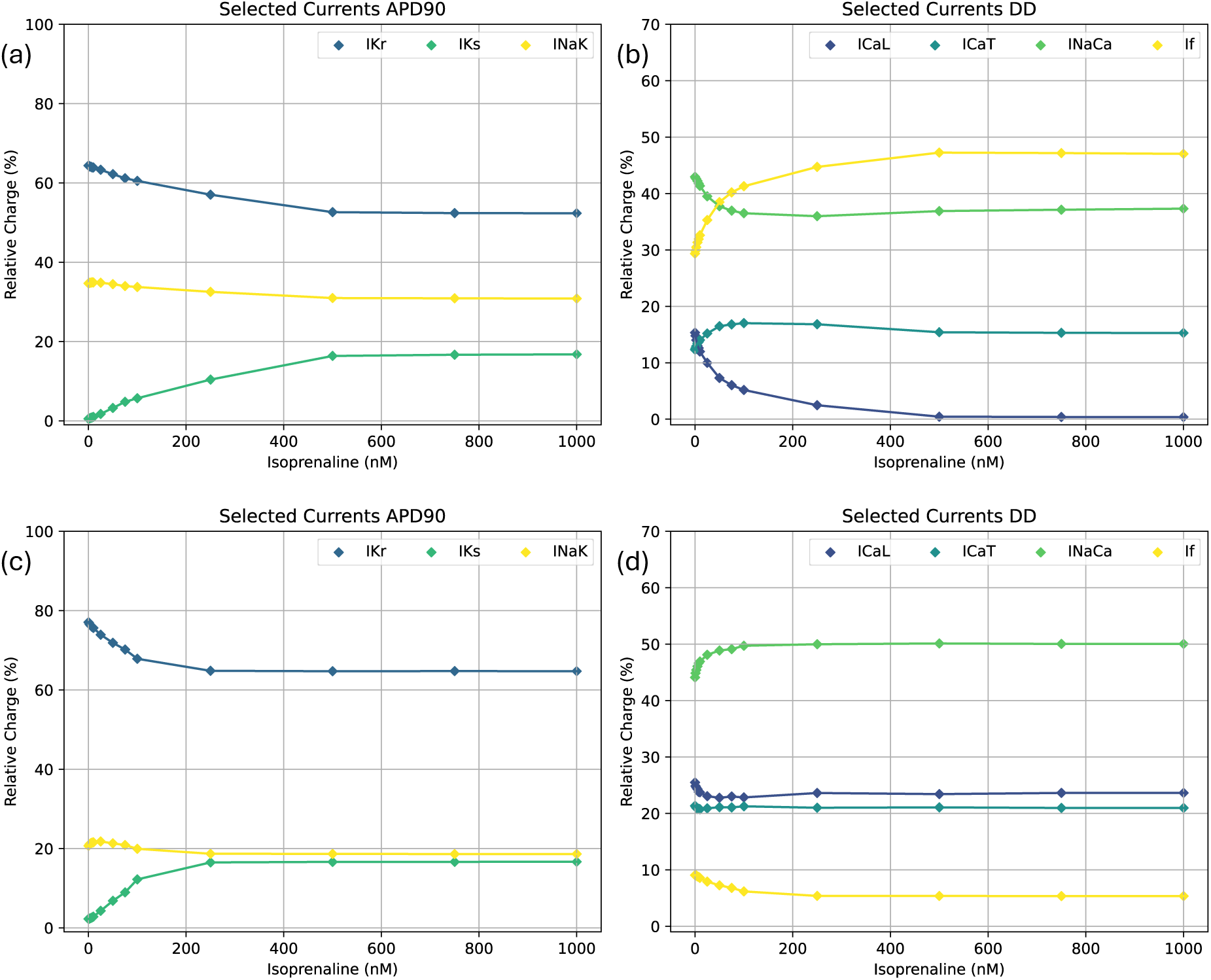
(a) relative contribution of selected currents to APDS0 (IKr in blue, INaK in yellow and IKs in green) and (b) DD (If in yellow, INaCa in light green, ICaL in blue, and ICaT in dark turquoise) for different [ISO] in the extended Severi model. (c) relative contribution of selected currents to APDS0 (IKr in blue, INaK in yellow and IKs in green) and (d) DD (If in yellow, INaCa in light green, ICaL in blue, and ICaT in dark turquoise) for different [ISO] in the extended Fabbri model. Note the different scales for the relative charge for APD90 and DD.

The contribution of I_CaL_ during DD was 0.3 and 1.8 pC, for the extended Severi and Fabbri model, respectively. For both models, the contribution decreased slightly at 10 nM [ISO] by 3.3 and 1.8 % (Figure 3b,d). Additionally, the charge via I_CaT_ and I_NaCa_ was assessed without being directly affected by the channel phosphorylation. When increasing [ISO] to 10 nM in the extended Severi model, the contribution of I_CaT_ with 0.3 pC increased by 1.7 %, while the impact of I_NaCa_ with 0.9 pC was reduced by 1.6 %. In contrast, in the extended Fabbri model, the contribution of I_CaT_ with 1.5 pC decreased by 0.6 % and the impact of I_NaCa_ with 3.1 pC was elevated by 2.8 %. In the extended Severi model, the more pronounced impact of I_f_ during basal conditions was further increased under β-AR stimulation. In the extended Fabbri model, the impact of I_f_ decreased with increasing [ISO], whereas I_NaCa_ transferred more charge. Although I_CaL_ lost relatively in significance considering the complete DD phase, the activation curve shift and increase in conductance led to an increased contribution in the early part of DD, increasing the depolarisation slope. These results were in accordance with the different emphasis on the membrane and Ca^2+^ clock in both models. However, the relatively small changes in current contributions could not fully explain the high inter-species differences observed in the BR increase under β-AR stimulation.

The comparison of further Ca^2+^-clock phosphorylation targets revealed high variations in the effect on RyR and SERCA, which control the sarcoplasmic Ca^2+^ release and uptake. For 10 nM [ISO] the maximum amplitude of Ca^2+^ release and uptake in the extended Severi model increased by 46.2 and 65.9 %, respectively, whereas the maxima in the extended Fabbri model increased by 159.9 and 275.6 %. This augmentation of the sarcoplasmic Ca^2+^ exchange resulted in higher I_NaCa_ activity both during the AP and DD extruding Ca^2+^ ions. The sarcoplasmic Ca^2+^ release by the RyR led to a largely increased cytoplasmic Ca^2+^ concentration during DD, which further activated the signalling cascade via calmodulin stimulated AC and activated I_CaL_ at earlier stages of the DD.

To analyse the currents during the AP, I_Kr_, I_Ks_ and I_NaK_, were integrated (Figure 3a,c). With increasing [ISO], the charge transferred by I_Ks_ increased caused by the conductance increase and activation curve shift, whereas the contribution of I_NaK_ and I_Kr_ decreased. 10 nM [ISO] in the extended Severi model yielded slightly elevated contributions of I_Ks_ with 0.03 pC and I_NaK_ with 1.7 pC (0.5 % and 0.2 % increase, respectively). The contribution of I_Kr_ with 3.2 pC decreased by 0.7 %. When further increasing [ISO] to 100 nM, the impact of I_Ks_ increased by 5.2 %, while the contribution of I_NaK_ and I_Kr_ decreased by 1.0 % and 4.2 %, respectively. Similarly, the contribution in the extended Fabbri model increased slightly for low [ISO], whereas 100 nM [ISO] led to an elevation of 10.0 % for I_Ks_ and reduction of 0.8 % and 9.2 % for I_NaK_ and I_Kr_, respectively. The absolute charge of I_Kr_, I_Ks_ and I_NaK_ increased slightly with elevated [ISO], while the decreased contribution of I_NaK_ and I_Kr_ was mainly counteracted by the largely increased transferred charge through I_Ks_. This led to the small prolongation of APD90 under β-AR stimulation.

### Effects of Hypocalcaemia

As previously reported, hypocalcaemia leads to increased CLs [4]. For a comparison of the underlying mechanisms responsible for the bradycardic effect, the extended Severi and Fabbri models were analysed for [Ca^2+^]_o_ gradually reduced from 1.8 mM (basal conditions) to 1.3 and 1.2 mM, respectively. Even lower [Ca^2+^]_o_ could occur in HD patients, but further reductions of [Ca^2+^]_o_ led to cessation of automaticity in both SANC models. The CL of the extended Severi model increased by a total of 39.4 % (343.0 to 478.2 ms) for 1.3 mM [Ca^2+^]_o_, while for the extended Fabbri model, the increase was 174.4 % (813.2 to 2231.8 ms) at 1.2 mM [Ca^2+^]_o_. DD was largely prolonged during hypocalcaemia, whereas the APD90 affected the CL prolongation only mildly in each of the models. Again, selected currents were integrated to assess the transferred charge during the different phases of the AP. For the extended Severi model, the contribution of I_f_, with a charge of 0.7 pC in the basal state, was increased by 13.8 % for 1.3 mM [Ca^2+^]_o_. Consequently, the Ca^2+^ cycling was attenuated and thus, the contribution of I_CaT_ and I_NaCa_ to membrane depolarisation was reduced. Additionally, the decreased Ca^2+^ influx attenuated the Ca^2+^ transient as well as the maximum amplitude of the SERCA uptake by 58.3 % and release via RyR by 50 %. Figure 4 shows the relative contribution to re- and depolarisation of selected currents for several [Ca^2+^]_o_. These results were in agreement with previous results of Kohajda et al., in which a decreased [Ca^2+^]_o_ reduced I_NaCa_ and lead to a more susceptible DD phase [1].

**Figure 4:**
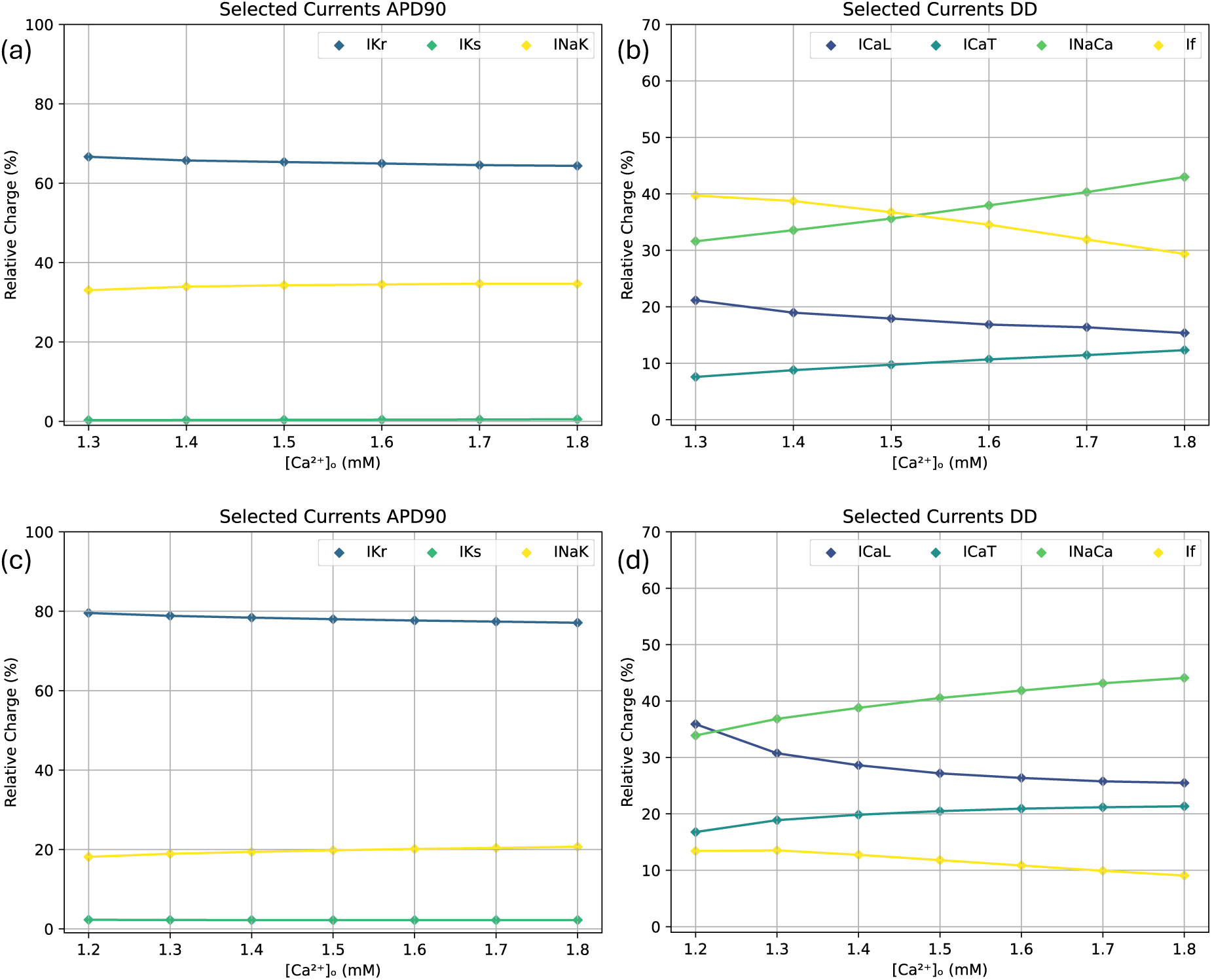
(a) relative contribution of selected currents to APDS0 (IKr in blue, INaK in yellow and IKs in green) and (b) DD (If in yellow, INaCa in light green, ICaL in blue, and ICaT in dark turquoise) for different [Ca^2+^]_o_ in the extended Severi model. (c) relative contribution of selected currents to APDS0 (IKr in blue, INaK in yellow and IKs in green) and (d) DD (If in yellow, INaCa in light green, ICaL in blue, and ICaT in dark turquoise) for different [Ca^2+^]_o_ in the extended Fabbri model. Note the different scales for the relative charge and the increasing [Ca^2+^]_o_ on the x-axis.

In contrast to the extended Severi model, I_CaL_, I_CaT_ and I_NaCa_ played more important roles in the extended Fabbri model. During early DD, I_CaT_ and I_NaCa_ dominated with 0.4 and 0.6 pC, equivalent to a share of 36.6 and 53.2 %, while I_f_ and I_CaL_ played a minor role. With decreasing [Ca^2+^]_o_, the impact of I_NaCa_ declined and I_f_ gained influence (Figure 4d). Less extracellular Ca^2+^ decreased the impact of Ca^2+^ in depolarising the membrane, thus the relative contribution of K^+^ and Na^+^ currents increased. In the basal state, the charge transferred during the complete DD mainly consisted of I_NaCa_ with 3.1 pC (44.3 %), I_CaL_ with 1.8 pC (25.3 %) and I_CaT_ 1.1 pC (21.4 %). Whereas for decreasing [Ca^2+^]_o_, the contribution of I_f_ and I_CaL_ was elevated by 4.3 and 10.5 %, respectively, the contribution of I_NaCa_ decreased by 10.0 % for 1.2 mM [Ca^2+^]_o_. The maximum amplitude of the SERCA uptake was attenuated by a factor of 10 and the Ca^2+^ release via RyR was reduced by 63.0 %.

In general, the depolarisation of the membrane depends on the influx and outflux of positive charged ions. In the SANC models, this could either be Ca^2+^ ions, which are exchanged between four different cell compartments (intra- and extracellular space, junctional and network SR) or K^+^ and Na^+^, which are exchanged between the intra- and extracellular space. The reduction of [Ca^2+^]_o_ resulted in slower Ca^2+^ influx through I_CaT_ and I_CaL_ during DD, which in turn led to a slower increase of cytoplasmic Ca^2+^ content and attenuated I_NaCa_ activity as well as Ca^2+^ uptake by SERCA and Ca^2+^ release by RyR. Meanwhile, repolarising currents during DD, like I_Kr_, I_Ks_ and I_NaK_ were attenuated to a lower extend, which prolonged the complete depolarisation process to reach TOP and increased I_f_ based on larger Na^+^ and K^+^ gradients. Consequently, the prolonged DD increased the absolute charge transferred during DD based on the decreased depolarisation slope, while the smaller Ca^2+^ uptake was compensated for by an increased I_NaCa_ activity during the APD. Thus, the relative contribution of Ca^2+^ ions to membrane depolarisation was slightly reduced and the relative contribution of I_f_, carrying K^+^ and Na^+^ ions, to reach TOP increased.

### Effects of Hypocalcaemia and Increased Sympathetic Tone

The influence of graded changes in [Ca^2+^]_o_ and [ISO] are shown in Figure 5. The BR increased largely for small [ISO], while the influence of [Ca^2+^]_o_ was rather linear for higher [ISO]. When dividing the AP cycle into APD90 and DD, in both extended models, a logarithmic-like increase with saturation could be observed for the DD (Figure 5c,f). For simulations with the extended Severi model with [Ca^2+^]_o_ near the basal conditions, the changes in the APD90 reached a maximum for rather small [ISO] (25 to 50 nM) (Figure 5b). Decreased [Ca^2+^]_o_ led to an attenuation of the maximal effect of [ISO]. For different [Ca^2+^]_o_, the BR changes in the extended Fabbri model remained nearly linear. While the DD was affected similarly, the peak of the APD90 for small [ISO] (25 to 50 nM) remained prominent, while further decreasing [Ca^2+^]_o_ extenuated the peak and the total APD prolongation (Figure 5e).

**Figure 5:**
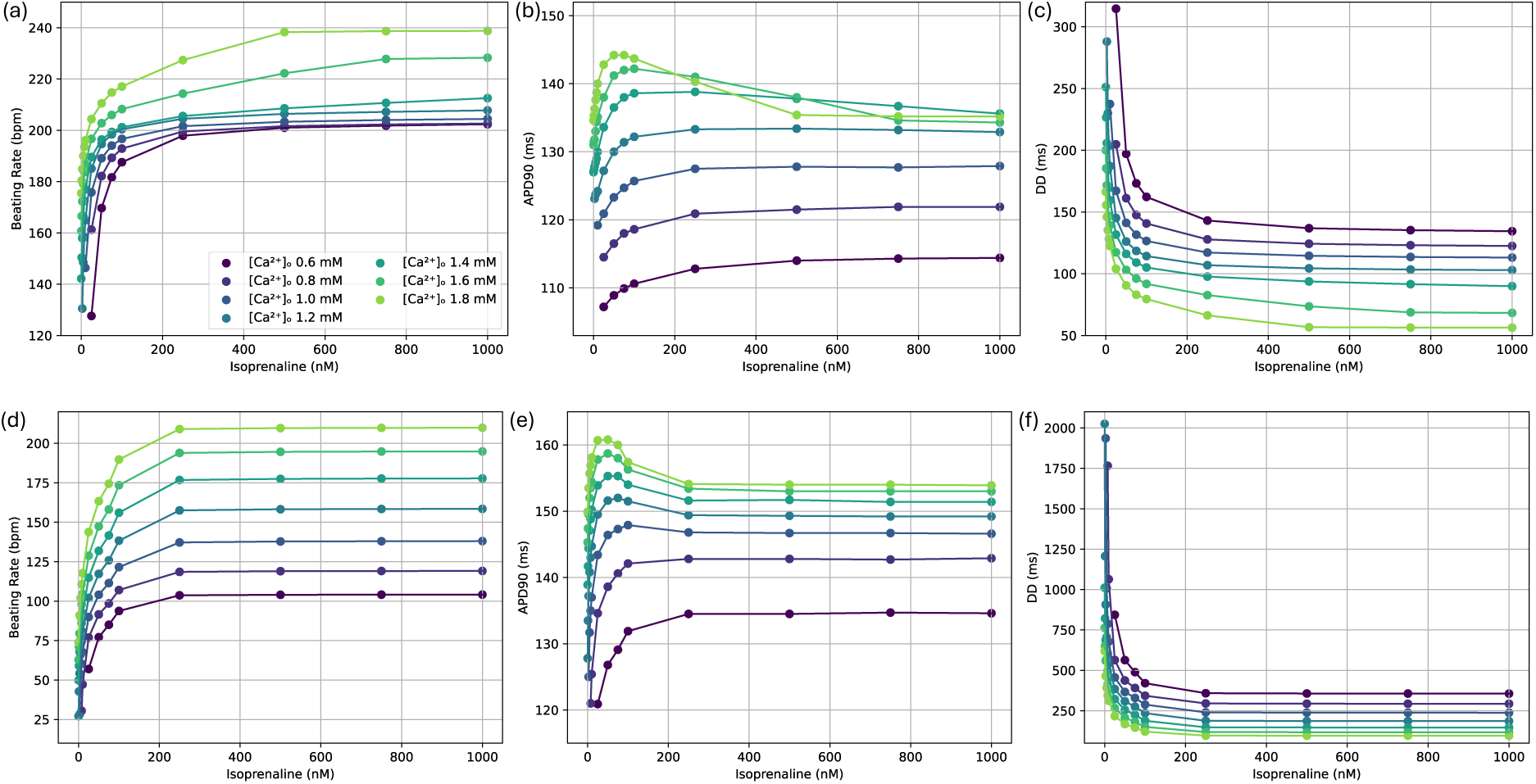
(a) BR, (b) APD90 and (c) DD for varying [ISO] and [Ca^2+^]_o_ in the extended Severi et al. (rabbit) model; (d) BR, (e) APD90 and (f) DD in the extended Fabbri et al. (human) model. Each line illustrates a different [Ca^2+^]_o_.

Analysing the compensatory effect of ISO during hypocalcaemia showed that for linear decreases in [Ca^2+^]_o_, an exponential-like increase in [ISO] was needed to maintain the basal BR. In the extended Severi model with [Ca^2+^]_o_ of 1.2 mM, 15.5 nM [ISO] restored the initial BR. Although ISO was applied and the basal BR restored, the charge transferred by the Ca^2+^ transient (CaT) remained smaller with a decrease of 38.2 % compared to basal conditions. Conversely, for the Fabbri model, 7.3 nM were sufficient, while the transferred charge of CaT decreased by 29.9 %. This highlighted that the extended Fabbri model is not only more sensitive to changes in [Ca^2+^]_o_ but even more so to sympathetic stimulation. Further decreasing [Ca^2+^]_o_ to a minimum of 0.6 mM required [ISO] of 60.0 nM and 41.7 nM, respectively, to maintain the basal BR. The charge transferred by CaT decreased by 72.7 and 61.9 %. Mimicking a sudden loss of sympathetic tone at low [Ca^2+^]_o_ ([ISO] dropping to 0 nM after 1000 s) unmasked the low basal BR under hypocalcaemic conditions. In the extended Severi model, this led to decreased BRs of 115 to 130 bpm before automaticity stopped within seconds. In contrast, the extended Fabbri model stabilised after a few seconds with extreme bradycardic BR of 10 to 35 bpm. Figure 6 illustrates the compensatory effect of ISO and the resulting BRs.

**Figure 6:**
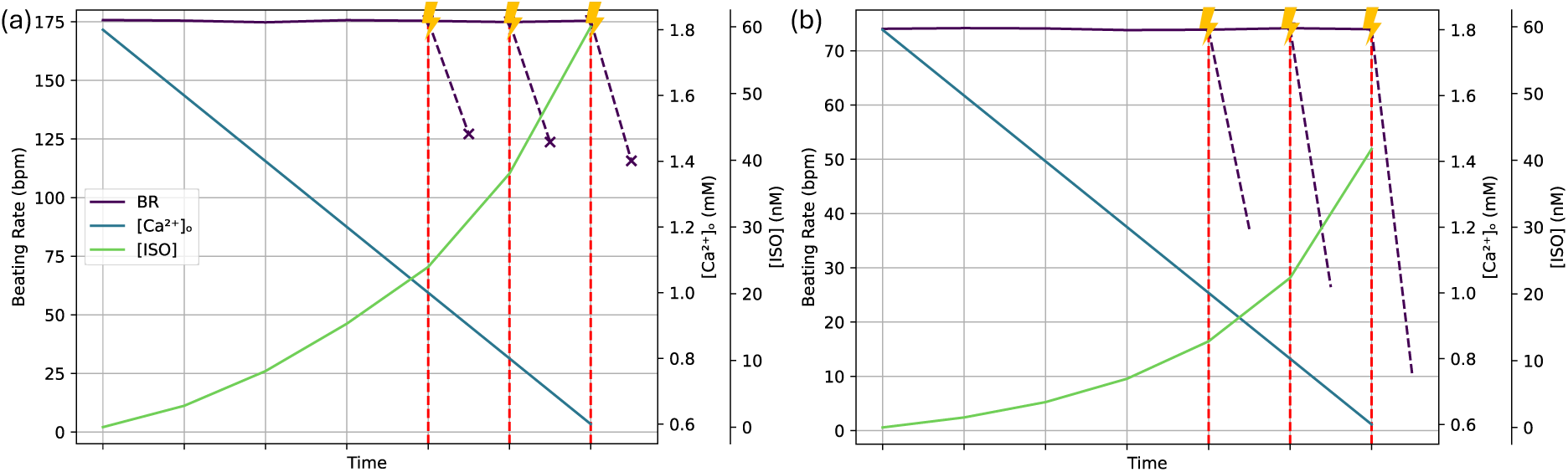
(a) compensatory effect of ISO in the extended Severi model; (b) in the extended Fabbri model. Decreasing [Ca^2+^]_o_ (dark turquoise) was compensated for by increased [ISO] (light green) to maintain the basal BR (purple). Flashes and vertical red dashed lines indicate three points in time at which [ISO] was set to 0 nM (sudden loss of sympathetic tone) after running the models for 1000 s to reach a steady CL. The purple dashed lines indicate the resulting BRs after a few seconds, x indicates cessation of automaticity.

Again, the AP cycle was divided into APD90 and DD to assess the charge of selected current contributions during these phases. Under basal conditions, APD90 was mainly determined by I_Kr_ and I_NaK_, in the extended Severi model. With increased [ISO], I_Ks_ gained influence due to the increase in conduction and activation curve shift, which led to an earlier and prolonged K^+^ ion flux duration into the extracellular space (Figure 7a). For decreasing [Ca^2+^]_o_ this effect was attenuated.

**Figure 7:**
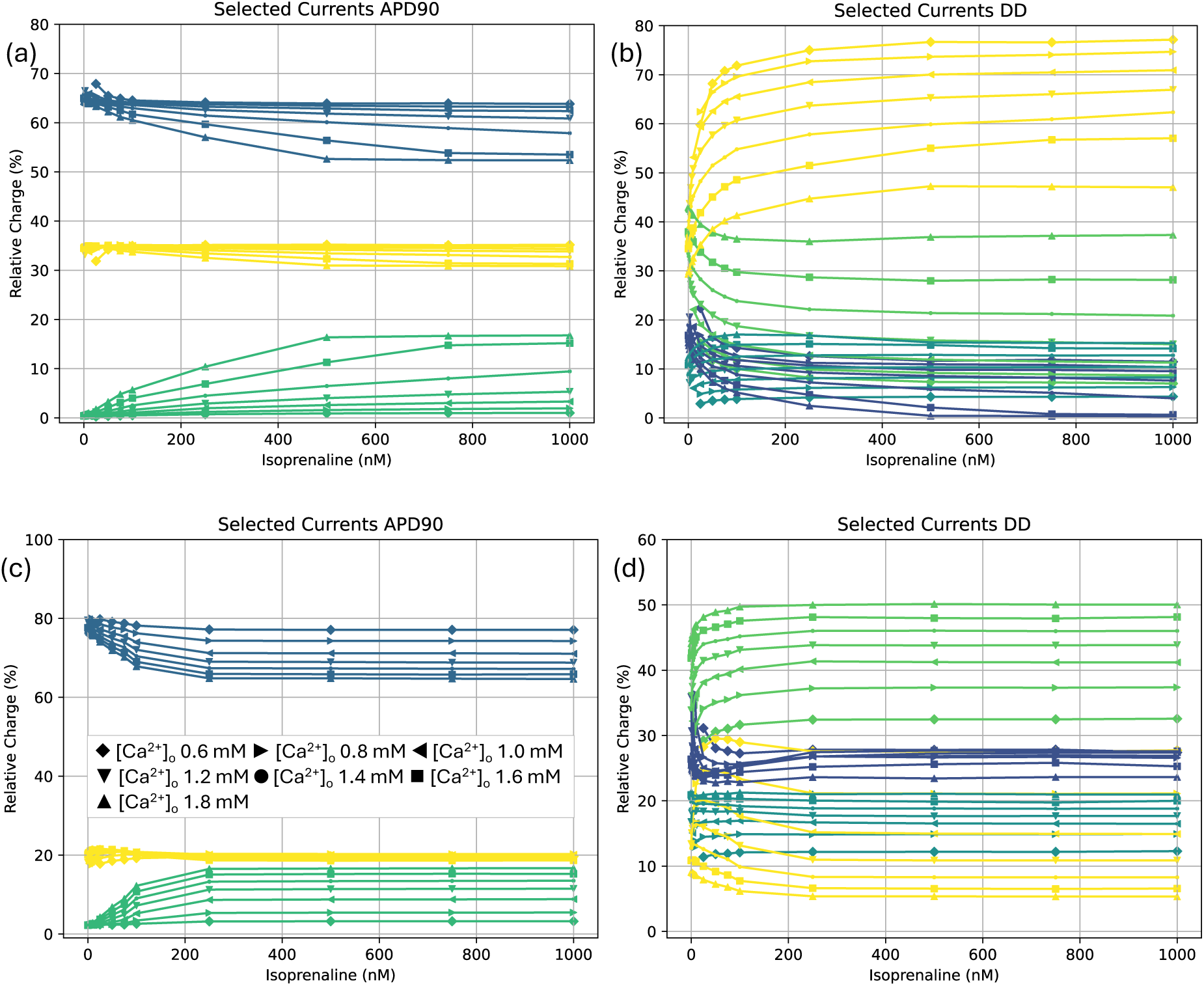
(a) relative contribution of selected currents to APDS0 (IKr in blue, INaK in yellow and IKs in green) and (b) DD (If in yellow, INaCa in light green, ICaL in blue, and ICaT in dark turquoise) for different [ISO] in the extended Severi model. (c) relative contribution of selected currents to APDS0 (IKr in blue, INaK in yellow and IKs in green) and (d) DD (If in yellow, INaCa in light green, ICaL in blue, and ICaT in dark turquoise) for different [ISO] in the extended Fabbri model. Each symbol represents a different [Ca^2+^]_o_. Note the different scales for the relative charge on the y-axis.

DD was dominated by I_NaCa_ with a contribution of 43.0 % and I_f_ with 29.4 %. I_CaT_ and I_CaL_ had smaller shares with 15.3 % and 12.3 %. When gradually increasing [ISO], the influence of I_f_ and I_CaT_ increased, while the contribution of I_NaCa_ and I_CaL_ decreased (Figure 7b). With decreasing [Ca^2+^]_o_ of 0.6 mM, I_f_ largely gained in significance. Whereas the contribution of I_CaL_ increased slightly, the influence of I_CaT_ and I_NaCa_ decreased. Consequently, the reduced cytoplasmic Ca^2+^ level attenuated the exchange between the cell compartments, which attenuated sarcoplasmic Ca^2+^ uptake and release. Thus, the increased conductance and activation curve shift by increased [ISO] on I_CaL_ was negligible.

Comparing these results to the extended Fabbri model, a similar but more pronounced trend for the APD90 could be observed: I_Kr_ dominated for baseline [Ca^2+^]_o_ of 1.8 mM and contributed less with increasing [ISO]. In contrast, I_Ks_ became more important for reduced [Ca^2+^]_o_ and increased [ISO] (Figure 7c).

DD (Figure 7d) was dominated by I_NaCa_, I_CaL_ and I_CaT_ with shares of 44.1 %, 25.5 % and 21.3 %, respectively. I_f_ contributed with only 9.1 %. With the gradual increase of [ISO], the influence of I_NaCa_ increased, whereas the impact of I_f_ and I_CaL_ decreased. The influence of I_CaT_ remained similar. Compared to the extended Severi model, the influence of the currents in the extended Fabbri model changed more linearly with respect to [Ca^2+^]_o_.

For both extended models, for increased [ISO], the membrane currents and Ca^2+^ cycling were more pronounced, which led to faster depolarisation. Thereby, the β-AR stimulation affected both clocks. In contrast, the depletion of [Ca^2+^]_o_ led to an attenuation of Ca^2+^ influx by I_CaL_ and I_CaT_, which was based on the smaller gradient between intra- and extracellular space. Consequently, the reduced cytoplasmic Ca^2+^ content reduced I_NaCa_ as well as the sarcoplasmic Ca^2+^ uptake and release, attenuating the Ca^2+^transient without directly altering the de- and repolarisation by K^+^ and Na^+^ ions. Thus, for decreased [Ca^2+^]_o_, the effect of increased [ISO] switched from ISO-induced increased Ca^2+^ cycling more towards K^+^ and Na^+^ exchange.

To assess the predominant mechanisms mediating sympathetic stimulation, numerical simulations, in which only one particular target was altered, while the other SR or membrane proteins were modelled as insensitive to PKA-phosphorylation and cAMP, were performed. In the extended Severi model, at the control [Ca^2+^]_o_ of 1.8 mM, the mostly affected targets for small [ISO] were I_f_, I_CaL_ and the Ca^2+^ uptake by SERCA. For 75 nM [ISO], the BR increased by 12.6 %, in case only the activation curve of I_f_ was shifted and by 12.7 %, in case of SERCA phosphorylation. Solely altering I_CaL_ increased the BR by 7.0 %. In comparison, modulating all targets of the ANS at 75 nM [ISO] elevated the BR by 25.7 %. For concentrations larger than 100 nM, SERCA was the most influential ion flux with a maximum BR increase of 32.1 % (Figure 8a, light green curve).

**Figure 8:**
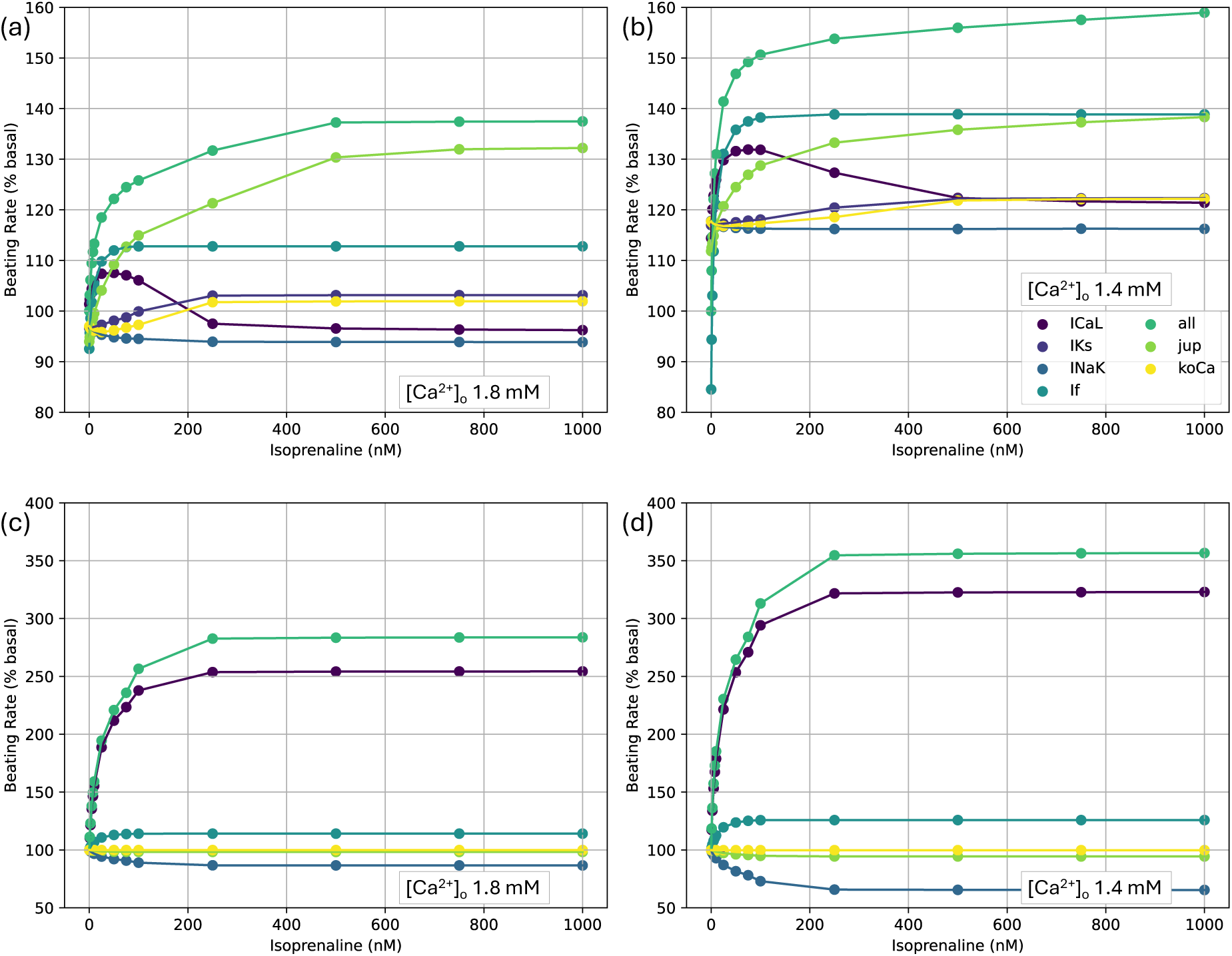
To assess the predominant mechanisms mediating sympathetic stimulation, numerical simulations, in which only one particular target was altered, while the other SR or membrane proteins were modelled as insensitive to PKA-phosphorylation and cAMP, were performed. Illustration of ISO affecting only one target for control [Ca^2+^]_o_ of 1.8 mM (left) and hypocalcaemia at 1.4 mM (right) with each altered target represented by a different colour. jup and koCa correspond to the SERCA uptake and a variable mediating the Ca^2+^ release by RyR, respectively. (a) extended Severi model for control [Ca^2+^]_o_ of 1.8 mM; (b) extended Severi model for [Ca^2+^]_o_ of 1.4 mM; (c) extended Fabbri model for control [Ca^2+^]_o_ of 1.8 mM; (d) extended Fabbri model for [Ca^2+^]_o_ of 1.4 mM.

Decreased [Ca^2+^]_o_ augmented the influence of β-AR stimulation. Lower [Ca^2+^]_o_ decreased the Ca^2+^ exchange between the cell compartments and, thus, I_f_ had a stronger influence on pacemaking. ISO only affecting I_f_ decreased the BR for small [ISO], whereas it increased the BR for higher [ISO]. This effect was more pronounced with lower [Ca^2+^]_o_. At 1.4 nM [Ca^2+^]_o_, the maximum BR increase for ISO only affecting I_f_ peaked at 38.9 % (Figure 8b, dark turquoise curve). In case only I_CaL_ was affected by phosphorylation, the impact on the BR peaked at about 50 nM [ISO], with an elevation of 31.6 % (Figure 8b, purple curve). Low [Ca^2+^]_o_ led to a more influential I_CaL_, whereas it attenuated the effect of the Ca^2+^ uptake (Figure 8b, light green curve). I_Ks_ and the sarcoplasmic Ca^2+^ release via RyR were more prominent for higher [ISO], still affecting the BR just slightly by 3.1 and 1.9 %. Since I_NaK_ was mainly active during APD, which marginally contributed to the CL shortening, the effects were negligible. Figure 8a,b summarise the simulation results of the extended Severi model, in which only one target was influenced by [ISO] in basal conditions and [Ca^2+^]_o_ of 1.4 mM.

In the extended Fabbri model, with control [Ca^2+^]_o_, I_CaL_ was most influential in increasing the BR during sympathetic stimulation. For [ISO] larger than 250 nM, only affecting I_CaL_ accelerated the BR by a maximum of 154.1 % (Figure 8c, purple curve). Solely affecting I_f_ also influenced pacemaking, peaking at only 14.2 % (Figure 8c, dark turquoise curve). For all other targets, the model predicted no significant changes in the BR.

When gradually decreasing [Ca^2+^]_o_, the influence of sympathetic stimulation on the BR became larger. For 1.4 mM [Ca^2+^]_o_, I_CaL_ remained the most important current, peaking at an increase of 222.9 % (Figure 8d, purple curve), while the influence of I_f_ increased to 72.6 % during hypocalcaemic conditions (Figure 8d, dark turquoise curve). Altering all targets led to a maximum increase of 256.6 %. Whereas the phosphorylation of the RyR was negligible, all other targets (I_Ks_, I_NaK_ and SERCA) reduced the BR. Due to the APD prolongation caused by I_Ks_ and I_NaK_, the model only yielded APs for [ISO] smaller than 50 nM and 5 nM, respectively, when considering only the alteration of these targets. This underlined the sensitive interaction between the components of the membrane and Ca^2+^ clock. Thus, increased ion fluxes during repolarisation had to be counteracted during depolarisation and vice versa. In case compensation was not possible due to asymmetrically high increases in one of the phases by the alteration of only one target, we observed increased CL or cessation of automaticity. Figure 8c,d illustrate the results of the extended Fabbri model under basal [Ca^2+^]_o_ of 1.8 mM and hypocalcaemia (1.4 mM), respectively.

### Model Validation

In basal conditions, the extended Severi model yielded a BR of 174.72 bpm. The BR was slightly higher than in the original Severi model (169.05 bpm) and slightly lower than in the BY model (182.70 bpm). All basal BRs were inside the physiological range (150 – 250 bpm) for fresh SAN rabbit cells [24]. In nearly all sources providing data for several ISO concentrations, elevated [ISO] increased the BR logarithmic. Considering [ISO] of 100 nM, the BR increased by 23.6 % in the extended Severi, as observed in the *in vitro* experiments [10].

Furthermore, [ISO] of 300 nM and 1000 nM resulted in BR elevations of 32.1 and 36.0 %, respectively. These predictions are in accordance with the experimental data by Vinogradova et al. [26]. Table 7 summarises the observed BRs in the extended Severi model, several experimental measurements on isolated rabbit SANCs and the effect of increased [ISO] on Langendorff-perfused hearts. The differences between isolated SANCs and measurements on Langendorff-perfused hearts might be caused by the enzymatic solutions used to dissociate the tissue to individual cells. Thereby, proteases naturally dissociated not only coupling structures, but also channels, receptors and possibly parts of the AC-cAMP-PKA signalling cascade, which led to reduced responses to b-AR stimulation. Nevertheless, the trend remained logarithmic, while the saturation might start at lower [ISO].

**Table 7:** Effects of ISO on the BR of isolated rabbit SANCs in experimental reports from literature and experiments on Langendorff-perfused hearts.

| [ISO] (nM) | Extended Severi | Isolated Single Cells (n=4,10) | Langendorff-perfused Hearts (n=5) |
| --- | --- | --- | --- |
| 0 | 174.72 bpm | (150.00 to 250.00) bpm [24] | 179.27 50.21) bpm |
| 10 | 195.44 bpm, + 11.7 % | (196.07 ± 65.88) bpm, + 7.1 % [14] | (248.88 ± 49.20) bpm, + 38.8 % |
| 20 | 201.61 bpm, + 15.3 % | (206.79 ± 88.50) bpm, + 13.0 % [14] | – |
| 50 | 209.79 bpm, + 19.9 % | – | (260.39 ± 85.46) bpm, + 45.3 % |
| 100 | 216.29 bpm, + 23.6 % | (223.05 ± 32.56) bpm, + 22.0 % [14] | (278.94 ± 70.14) bpm, + 55.6 % |
| 300 | 230.77 bpm, + 32.1 % | (228.40 ± 21.66) bpm, + 31.7 % [26] | – |
| 500 | 237.34 bpm, + 35.7 % | – | (293.80 ± 18.75) bpm, + 63.9 % |
| 1000 | 237.91 bpm, + 36.0 % | (235.77 ± 61.13) bpm, + 35.5 % [26] | – |

The PKA level for varying β-AR stimulation in both extended models ranged from 0.72 to 0.97, similar to experimental values and simulations with the BY model [14]. The maximum of the introduced cAMP level in the extended Severi model was lower than in the original BY model 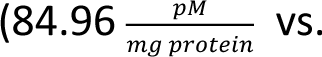 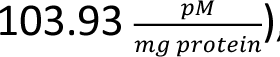, while the extended Fabbri model exhibited similar cAMP concentrations 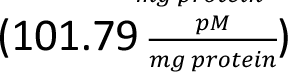.

## Discussion

In this study, the inter-species differences in the SANC response to hypocalcaemia [15], β-AR stimulation and the combination of both mechanisms, i.e. the compensation for hypocalcaemia-induced bradycardia by the ANS, were analysed in extended computational models of rabbit (Severi) and human (Fabbri) SANCs. The analysis of the results revealed that: (1) In the extended Severi model, the primary pacemaker currents during DD are I_f_ and I_NaCa_, whereas for the extended Fabbri model, pacemaking is rebalanced towards I_CaT_, I_CaL_ and I_NaCa_. (2) The integration of the β-AR signalling cascade increases Ca^2+^ inflow and cycling during DD (I_CaL_, SR Ca^2+^ uptake and release directly and I_CaT_, I_NaCa_ indirectly) and thus shortens DD and CL. (3) A reduction of [Ca^2+^]_o_ results in decreased Ca^2+^ inflow and cycling (I_CaT_, I_CaL_, SR Ca^2+^ uptake and release), prolonging DD and CL. (4) Hypocalcaemia is compensated for by an increased sympathetic tone, since similar targets were affect by both mechanisms. (5) The extended Fabbri model is more sensitive to sympathetic stimulation and hypocalcaemia as transmembrane depolarisation by Ca^2+^ is more accentuated.

### Inter-Species Differences Under Basal Conditions

In the past, there was different emphasis on the role of Ca^2+^ cycling vs. the contribution of I_f_. Despite a detailed description of intracellular Ca^2+^ cycling, I_f_ and I_NaCa_ were the primary pacemaker currents in the original Severi model. In the original Fabbri model, the impact of I_f_ was attenuated, which rebalanced the pacemaking control more towards I_CaT,_ I_CaL_ and I_NaCa_. This remained similar in the extended models. Thus, in the extended Fabbri model, the reduced contribution of I_f_ (compared to the extended Severi model) during early DD was primarily compensated for by the increased impact of I_CaT_. During late DD the compensation for the decreased contribution of I_f_ shifted towards I_CaL_, with the charge transferred by I_CaT_ declining due to channel inactivation. Since the activation of I_CaL_ was controlled by voltage and Ca^2+^-activated gates, most charge was transferred shortly after opening the Ca^2+^ gate by an increase in cytoplasmic Ca^2+^ concentration, which was caused by sarcoplasmic Ca^2+^ release from RyR. Delayed sarcoplasmic Ca^2+^ release attenuated the slope of I_CaL_ and delayed the rapid downstroke of a major contributor to reaching TOP and initiating the AP upstroke. Since repolarizing currents (e.g., I_Kr_, I_Ks_, and I_NaK_) were similarly active in both models, the different contribution during DD led to a prolonged depolarisation phase and greater total charge transferred during DD compared to the extended Severi model. As a result, in the extended Fabbri model, the impact of Ca^2+^ cycling was amplified, the CL was prolonged, and thus the BR was reduced.

### Logarithmic Beating Rate Increase Under Sympathetic Stimulation

By integrating the β-AR signalling cascade of Behar et al. into both models, continuous modelling of β-AR effects on critical membrane (I_CaL_, I_f_, I_Ks_, I_NaK_) and SR (Ca^2+^ release by RyR and uptake by SERCA) currents was enabled. Increases in [ISO] led to an increased cAMP and PKA activity level, which affected the coupled clock mechanisms, elevating the BR [14]. For increased ISO larger than 500 and 250 nM, for the extended Severi and Fabbri model, respectively, a saturation effect was observed. This was consistent with the implementation of the conductance increases and activation curve shifts as Hill curves.

The comparison of sympathetic stimulation affecting the BY and extended Severi rabbit SANC model unveiled that the extended Severi model predicted a stronger maximal response to β-AR stimulation on the BR (+36.0 %) than the BY model (+22.0 %). While the BY model closely reproduced the experimental data of cultured SANCs conducted by Yaniv et al. [19], several studies implied larger BR increases and slower saturation effects with respect to autonomic stimulation by elevated [ISO] [25], [26], [27], [23]. Yang et al. analysed the differences between fresh and cultured rabbit SANCs. The results suggested that cultured SANCs underlay a partial disengagement of the coupled clock mechanisms. While the reduction in PKA-dependent phosphorylation of PLB and RyR was partly restored by increased β-AR stimulation (1 nM [ISO]), the phosphorylation of further critical surface membrane channels, such as I_CaL_, I_Ks_ or I_NaK_ was not assessed and might also be affected [28]. Since the parametrisation of the BY model was based on cells cultured for 48 hours, with 1 nM [ISO] referring to basal conditions to counteract the decreased BR, the maximal effect of the ANS might also be attenuated. The logarithmic BR increase in combination with the restoration of the uncoupling effects of PLB and RyR could explain the discrepancy between the experimental data of cultured and fresh SANC and the different BR increases of the BY model compared to the extended Severi model. Additional differences between both rabbit models were mainly based on the different [Ca^2+^]_o_ used as basal conditions (2.0 mM BY vs. 1.8 mM Severi model) and differently weighted contributions of the membrane and Ca^2+^ clock. Alike its parental model, the extended Severi model emphasises I_f_ as a main contributor of the membrane clock, whereas the BY model focuses on a more pronounced Ca^2+^ cycling.

It could be further argued that even the maximum BR increase of 36.0 % in the extended Severi model was still an underestimation, when considering the maximum BR increase of 63.9 % in the Langendorff-perfused hearts for 500 nM [ISO]. This might be caused by the isolation process and thus, the reduced number of channels, receptors or the uncoupling of β-AR signalling cascade components, which attenuated the β-AR cell response. Nevertheless, the logarithmic BR increases with saturation were observed for single cell and Langendorff-perfused heart experiments. Furthermore, Fabbri et al. investigated the propagation of human SAN APs into the surrounding atrial tissue, by an *in silico* study, with an 1D strand of homogeneously coupled SAN and atrial elements. They showed that the load of the atrial tissue, due to an increased contribution of I_Na_, was responsible for a faster upstroke of the SAN elements close to the atrial ones. Even if the two discrete coupled regions and a 1D strand, are far from a physiological configuration, the electrical load might also increase the BR *in vivo* [29].

### Target Specific Effects Under Sympathetic Stimulation

By ISO only modulating I_f_, in the extended Severi model, the BR increased to a maximum of 12.6 %. This highlighted that I_f_ plays a dominant role in mediating the chronotropic effect under autonomic regulation. Thereby, I_f_ increased the steepness of the DD slope near the MDP, which shortened the depolarisation of the membrane during early DD. In contrast to affecting one of the other targets, only the phosphorylation of SERCA was more pronounced for high [ISO], with a maximum BR elevation of 32.1 %. This increased the Ca^2+^ uptake and resulted in an increased sarcoplasmic Ca^2+^ release by RyR, which increased I_CaL_ and I_NaCa_. Thus, depolarisation slope was steeper, which shortened DD significantly.

Solely augmenting I_CaL_ led to increased BRs for small and the inverse effect for high [ISO]. The modulation of I_CaL_ resulted in a shortened late phase of the DD and increased upstroke velocity of the AP. The inverted effect was based on larger Ca^2+^ ion transport through an increase in conductance and activation curve shift. Thus, the time during which positively charged ions entered the cytoplasm during late DD and the early stages of the AP upstroke were prolonged. Without subsequent elevations in SERCA and I_NaCa_ activity, a largely increased Ca^2+^ influx prolonged the repolarisation and, thus, the CL.

The phosphorylation of the sarcoplasmic Ca^2+^ release by RyR prolonged the CL for small [ISO], while larger [ISO] slightly increased the BR. The mild influence of the phosphorylation-induced change in RyR, which resulted in an increase in sarcoplasmic Ca^2+^ release, was disproportionately affected by the unchanged SERCA uptake, which was adequately represented in the extended Severi model [30].

Similarly, I_Ks_ elevated the BR for increasing [ISO]. By shifting the voltage-dependent activation curve, the K^+^ ion channel gates opened earlier in time and at lower voltages, which, in combination with the increased maximum current amplitude, led to a faster repolarisation and moderate shortening of the APD. Instead, solely introducing conductance increases in I_NaK_, yielded APD prolongations, which slightly increased the AP CL. Compared to I_Ks_ this meant that the heightened K^+^ influx without the ion channel opening earlier in time based on an activation curve shift, led to a prolonged time of K^+^ extrusion, which increased the APD.

Based on the lack of experimental data, the PKA-mediated modulation of I_Ks_ and I_NaK_ was not included in the BY model implementation. However, omitting the influence of I_Ks_ in the extended Severi model led to BRs peaking at 100 nM [ISO]. Besides, when [ISO] was further increased, the BR decreased, which was caused by increased APD and AP upstroke. The influence of I_f_ in the extended Severi model in combination with the cAMP-induced activation gate shift without subsequent modulations in I_Ks_, extruding K^+^ ions during the APD resulted in non-physiological prolongations. For the sake of completeness, the changes on I_NaK_ were also included.

In the extended Fabbri model, in which pacemaking was rebalanced towards I_CaT_ and I_CaL_, only modulating I_CaL_ with the other β-AR targets remaining unchanged largely influenced the BR. Since, Ca^2+^ cycling between the cell compartments was more pronounced in basal conditions, SERCA and I_NaCa_ were able to extrude the increased amount of Ca^2+^ ions under ANS stimulation, without prolonging the CL as it was the case in the extended Severi model.

In case only I_f_ was modulated, the activation curve shift resulted in earlier opening of the channel, but the decreased impact of I_f_ influenced the BR acceleration less pronounced. With decreasing [Ca^2+^]_o_, the contribution of I_f_ increased, which was caused by the relatively elevated significance of K^+^ and Na^+^ fluxes in depolarising the membrane, whereas the Ca^2+^ cycling lost in significance. The reduced [Ca^2+^]_o_, attenuated I_CaT_ and I_NaCa_, and thus, the cytoplasmic Ca^2+^ content, which subsequently attenuated the sarcoplasmic Ca^2+^ uptake and release by RyR.

While at [Ca^2+^]_o_ near the basal conditions, solely augmenting I_Ks_ had negligible effects, the alteration of I_NaK_ slightly elevated the BR. At 1.2 mM [Ca^2+^]_o_, the BR decreased and prevented AP generation for higher [ISO]. In both cases, this originated from the elevated K^+^ and Na^+^ exchange between the intra- and extracellular space at low [Ca^2+^]_o_. Without simultaneous counteraction, APD was prolonged and repolarisation of the transmembrane prevented.

Similarly, in the basal state modulating only the SERCA uptake resulted in negligible BR changes, while for reduced [Ca^2+^]_o_ the CL was prolonged slightly. Under extreme hypocalcaemic conditions, the reduced Ca^2+^ uptake inhibited the release by the RyR and the activation of I_CaL_ and I_NaCa_ caused by the reduced [Ca^2+^]_i_. Thus, the DD was prolonged, initiating the AP upstroke later. Again, this highlighted the critical role of the SERCA uptake inhibiting the Ca^2+^ release by the RyR and the tightly controlled Ca^2+^ balance of the different compartments influencing each other [30].

Contrary to the extended Severi model, the attenuation of I_f_ resulted in reduced K^+^ and Na^+^ ion exchanges between the extra- and intracellular space, which also extenuated K^+^ and Na^+^ ion transfer to maintain homeostasis via I_Ks_ and I_NaK_. Instead, the Fabbri model was based on a more pronounced Ca^2+^ exchange between the different cell compartments, especially Ca^2+^ influx from the extracellular space by I_CaT_ and I_CaL_. Consequently, the sarcoplasmic Ca^2+^ uptake by SERCA and release by RyR were elevated. In combination with the increases in Ca^2+^ cycling components caused by phosphorylation, this led to higher sensitivity with respect to β-AR stimulation.

The simultaneous occurrence of cAMP-mediated gate shifting along with phosphorylation of the membrane and SR targets exerted the largest effect on the BR in both extended SANC models. To achieve a reduction of the entire CL, the elevated ion exchanges during depolarisation needed to be compensated for by increased currents during repolarisation and vice versa. Thus, the intricately orchestrated interplay by the membrane and Ca^2+^ clock could increase the resulting BR. This involved an earlier initiation and steeper DD slope, facilitated by the activation curve shift of I_f_, coupled with increased Ca^2+^ cycling through augmented I_CaL_, sarcoplasmic Ca^2+^ uptake and release. Consequently, the DD phase was rapidly shortened, whereas the APD remained akin to the basal conditions. A prolongation of the APD was prevented by the increased repolarisation currents I_Ks_ and I_NaK_.

### Hypocalcaemia-Induced Beating Rate Reduction

Loewe et al. analysed the effects of altered electrolyte levels on a slightly modified human SANC model of Fabbri et al. [4]. The results indicated that hypocalcaemia markedly impaired pacemaking, while alterations in the extracellular K^+^ and Na^+^ levels yielded negligible and mild effects on the BR, respectively. Thereby, hypocalcaemia-induced slowing of the sinus node pacemaking increased the total CL for both rabbit and human SANC models. The reduction of [Ca^2+^]_o_ attenuated the amount of Ca^2+^ ions entering the cell from the extracellular space, which also reduced the intracellular Ca^2+^ cycling. Accordingly, I_CaT_, I_CaL,_ I_NaCa_, the Ca^2+^ uptake by SERCA and release by RyR were decreased, while the contribution to the depolarisation of I_f_ was increased. Caused by the larger impact of I_f_, pacemaking in the extended Severi model was less sensitive regarding hypocalcaemia, which resulted in smaller CL increases compared to the extended Fabbri model.

### An Increased Sympathetic Tone Compensates for Hypocalcaemia

Assessing the compensatory effect of the ANS with respect to hypocalcaemia-induced bradycardia, typically occurring in HD patients, both rabbit and human models predicted that reduced [Ca^2+^]_o_ can be compensated for to a certain extent by an increased sympathetic tone. Stary et al. implemented a linear [ISO] dependence in the Fabbri model, which affected the BR almost linearly [8]. Contrary to these findings, the integration of the non-linear β-AR signalling cascade led to a logarithmic increase with saturation of the BR. This was similarly observed in several experiments on isolated rabbit SANCs [25], [26], [27], [23] as well as the Langendroff-perfused heart experiments. Furthermore, this was in accordance with the continuous implementation of the sympathetic effects, on critical membrane and SR currents, as Hill functions. Thus, the stimulation modulated the targets nearly linearly close to the basal conditions, with a saturation effect for more pronounced changes. The combination of both, hypocalcaemia, which attenuated the Ca^2+^ cycling, and the β-AR stimulation, which increased the influence of six different targets (I_CaL_, I_f_, I_NaK_, I_Ks_, RyR, SERCA), revealed that three of these targets were directly affected by [Ca^2+^]_o_ depletion and increased conductance and activation curve shifts by the ANS. Thus, hypocalcaemia-induced attenuations of the Ca^2+^ cycling were directly neutralised by an increased sympathetic tone. With less Ca^2+^ available, the increasing effects needed to be more pronounced to restore the basal BR. The other three modulated β-AR targets controlled the exchange of K^+^ and Na^+^ of the intra- and extracellular space. The combination of the attenuated I_f_ in the extended Fabbri model, with the increased impact of the Ca^2+^ clock components in regulating the BR explained the higher sensitivity of the extended Fabbri model towards hypocalcaemia and β-AR stimulation.

Verkerk et al. discussed changes in the kinetics of I_Ks_ with respect to β-AR stimulation in human SANC [31]. Based on the experimental data, using HEK-293 cells, the K^+^:Na^+^ permeability, conductance, and reversal potential were adjusted. Including these findings into the extended Fabbri model reduced the basal BR slightly to 70.41 bpm and the ISO-induced BR acceleration to a maximum of 177.78 bpm (+152.5 %). It can be argued that the increased BR of 177.78 bpm is closer to the physiologically achievable maximum of an adult human than the 209.21 bpm encountered using the I_Ks_ formulation of the original Fabbri model. Additionally, the complete integration of the cholinergic receptor stimulation could further enhance the extended models towards more realistic predictions. This could include simulations with a sudden increase in the parasympathetic tone.

### Limitations

During the integration process of the β-AR signalling cascade into the SANC models, some assumptions were made due to scarcity of experimental data. Since, the generation and degradation of intracellular cAMP are influenced by several model parameters, these equations were adjusted to cover the inter-species differences. Besides, the PKA-dependent activation curve shift of I_CaL_ and I_Ks_ was expected to behave similar to the cAMP-mediated activation curve shift of I_f_. Lastly, the increase in conductance for I_NaK_ and I_Ks_ were assumed to be similar to the PKA-mediated phosphorylation of I_CaL_. To ensure a more realistic sympathetic stimulation, channel specific differences in reference to changes in the PKA level should be further investigated.

Loewe et al. extended the Fabbri model to simulate variable [K^+^]_i_ and [Na^+^]_i_ with a constant basal sympathetic tone [4]. These optimisations also resulted in more stability towards lower [Ca^2+^]_o_. Further decreases of the Ca^2+^ levels in combination with lower [ISO] as well as variable K^+^/Na^+^ formulations might also be of interest to uncover the pathogenesis of hypocalcaemia-induced bradycardia. The original and extended Severi model implementation contained variable [Na^+^]_i._ However, fixing [Na^+^]_i_ to 7.5 mM (initial value) to ensure similarity of [Na^+^]_i_ formulation altered the simulation results negligibly. The effect of ISO increased marginally for small and decreased for high [ISO]. To improve the models towards more realistic effects of changes in electrolyte levels, variable [K^+^]_i_ and [Na^+^]_i_ should be implemented in both extended models.

Moreover, automaticity was only assessed on single cell level. To capture the excitation initiating the heartbeat *in vivo*, complementing the simulations on single cell level with tissue patches of coupled SANC and including the surrounding myocardium are necessary [32]. Furthermore, the experimental results of the Langendorff-perfused hearts indicated higher sensitivity towards increased [ISO] compared to single cell experiments. However, the modulations of specific membrane and SR channels are modelled based on isolated single cell experiments and thus, the resulting conductance increases and activation curve shifts, should be complemented by experiments with Langendorff-perfused hearts.

It is also worth to mention that both SANC models by Severi et al. and by Fabbri et al. were derived based on patch-clamp experiments on isolated single cells in Tyrode’s solution. Unfortunately, as Severi et al. pointed out, the [Ca^2+^]_o_ of standard Tyrode’s solution (1.8 or 2.0 mM) is significantly far from physiological Ca^2+^ concentrations in blood serum (1.0 to 1.3 mM) and, consequently, in the interstitial fluid, which represents the *in vivo* extracellular milieu. As far as the aim is a comparison of *in vitro* experimental data with the simulated electrical activity of cardiac cells, it is obviously correct imposing the same extracellular concentration used in experimental protocols. However, the usage of the same concentration can be incorrect, when the analysis of *in vivo* pathophysiological mechanisms is the ultimate aim of simulation, as it was the case for simulated hypocalcaemia and the compensatory effect by an increased sympathetic tone [33].

## Conclusion

In this computational study, inter-species differences between the extended Severi (rabbit) and Fabbri (human) SANC models were investigated to elucidate the pathophysiological mechanisms of the elevated prevalence of SCD in HD patients. Furthermore, we formulated and experimentally tested the hypothesis that increased sympathetic tone may compensate to some extent for hypocalcaemia-induced bradycardia, whereas sudden loss under hypocalcaemic conditions could lead to severe bradycardia and failure of the sinus node. By affecting the Ca^2+^ clock, extracellular Ca^2+^ depletion was balanced by β-AR stimulation of critical membrane and SR currents. While the Ca^2+^ cycling was directly affected by hypocalcaemia and the increased sympathetic tone, K^+^ and Na^+^ exchange played a subsidiary role, especially in the human-based SANC model. These findings could help to further understand the underlying pathomeachanisms of SCD in CKD patients. Furthermore, regular non-invasive point-of-care monitoring of electrolyte levels, especially [Ca^2+^]_o_, by electrocardiograms could allow early diagnosis and continuous or retrospective assessment of plasma electrolyte concentrations [34]. This might be a possibility to reduce the high prevalence of SCD in CKD patients in the future.

## Supporting information

Supplementary Material - Equations

## Acknowledgments

This work was supported by DFG-507828355 (CARPe-diem) and by the European High-Performance Computing Joint Undertaking EuroHPC under grant agreement No 955495 (MICROCARD) co-funded by the Horizon 2020 programme of the European Union (EU), the French National Research Agency ANR, the German Federal Ministry of Education and Research, the Italian ministry of economic development, the Swiss State Secretariat for Education, Research and Innovation, the Austrian Research Promotion Agency FFG, and the Research Council of Norway, and NKFIH-OTKA FK-142949, and the National Academy of Scientist Education.

## Data Availability

All data supporting the findings of this study are available within the paper or supplementary material.

## Author Contribution

M.L.: conception and design, analysis and interpretation of data, drafting, final approvement of the manuscript, agreement to be accountable for all aspects of the work; T.S. analysis and interpretation of data, revising critically for important intellectual content, final approval of manuscript, agreement to be accountable for all aspects of the work; G.B.: acquisition of data, drafting, final approval of manuscript, agreement to be accountable for all aspects of the work; N.N.: acquisition of data, analysis and interpretation of data, revising critically for important intellectual content, final approval of manuscript, agreement to be accountable for all aspects of the work; A.L.: conception and design, analysis and interpretation of data, revising it critically for important intellectual content, final approval of manuscript, agreement to be accountable for all aspects of the work.

