## Supplementary Material - Equations for "Sympathetic Stimulation Can Compensate for Hypocalcaemia-Induced Bradycardia in Human and Rabbit Sinoatrial Node Cells"

#### 1 Model Parameters

##### 1.1 Cell Compartments

| | Severi model | Fabbri model | Description |
|---|---|---|---|
| $Control$ | $32\,pF$ | $57\,pF$ | Cell electric capacitance |
| $L_{cell}$ | $70\,\mu m$ | $67\,\mu m$ | Cell length |
| $R_{cell}$ | $4\,\mu m$ | $3.9\,\mu m$ | Cell radius |
| $L_{sub}$ | $0.02\,\mu m$ | $0.02\,\mu m$ | Distance between jSR and surface membrane (submembrane space) |
| $V_{i,part}$ | $0.46$ | $0.46$ | Part of cell volume occupied with myoplasm |
| $V_{jsr,part}$ | $0.0012$ | $0.0012$ | Part of cell volume occupied by jSR |
| $V_{nsr,part}$ | $0.0116$ | $0.0116$ | Part of cell volume occupied by nSR |
| $V_{cell}$ | $\pi \cdot R_{cell}^2 \cdot L_{cell}$ | $\pi \cdot R_{cell}^2 \cdot L_{cell}$ | Cell volume |
| $V_{sub}$ | $2 \cdot \pi \cdot L_{sub} \cdot (R_{cell} - \frac{L_{sub}}{2}) \cdot L_{cell}$ | $2 \cdot \pi \cdot L_{sub} \cdot (R_{cell} - \frac{L_{sub}}{2}) \cdot L_{cell}$ | Submembrane space volume |
| $V_i$ | $V_{i,part} \cdot V_{cell} - V_{sub}$ | $V_{i,part} \cdot V_{cell} - V_{sub}$ | Myoplasmic volume |
| $V_{jSR}$ | $V_{jSR,part} \cdot V_{cell}$ | $V_{jSR,part} \cdot V_{cell}$ | Volume of jSR ($Ca^{2+}$ release store) |
| $V_{nSR}$ | $V_{nSR,part} \cdot V_{cell}$ | $V_{nSR,part} \cdot V_{cell}$ | Volume of nSR ($Ca^{2+}$ uptake store) |

##### 1.2 Fixed Ion Concentrations

| | Severi model | Fabbri model | Description |
|---|---|---|---|
| $Cao$ | $1.8\,\mu M$ | $1.8\,\mu M$ | Extracellular $Ca^{2+}$ concentration |
| $Ki$ | $140\,\mu M$ | $140\,\mu M$ | Intracellular $K^+$ concentration |
| $Ko$ | $5.4\,\mu M$ | $5.4\,\mu M$ | Extracellular $K^+$ concentration |
| $Nao$ | $140\,\mu M$ | $140\,\mu M$ | Extracellular $Na^+$ concentration |
| $Nai$ | $variable$ | $5.0\,\mu M$ | Intracellular $Na^+$ concentration |
| $Mgi$ | $2.5\,\mu M$ | $2.5\,\mu M$ | Intracellular $Mg^{2+}$ concentration |

#### 1.3 Variable Ion Concentrations

| | Description |
|---|---|
| $Cai$ | Intracellular $Ca^{2+}$ concentration |
| $Ca_{jsr}$ | $Ca^{2+}$ concentration in the jSR |
| $Ca_{nsr}$ | $Ca^{2+}$ concentration in the nSR |
| $Ca_{sub}$ | $Ca^{2+}$ concentration in the subspace |
| $Nai$ | Intracellular $Na^{+}$ concentration (constant in Fabbri et al. model) |

#### 1.4 Ionic Values

| | Severi model | Fabbri model | Description |
|---|---|---|---|
| $F$ | $96485.3415\,\frac{C}{M}$ | $96485.0\,\frac{C}{M}$ | Faraday constant |
| $R$ | $8314.472\,\frac{J}{kM \cdot K}$ | $8314.472\,\frac{J}{kM \cdot K}$ | Universal gas constant |
| $T$ | $310\,K$ | $310\,K$ | Absolute temperature for $37\,^{\circ}C$ |
| $RTONF$ | $\frac{R \cdot T}{F}$ | $\frac{R \cdot T}{F}$ | factor = $26.72655\,mV$ |
| $E_{Na}$ | $RTONF \cdot log(\frac{Nao}{Nai})$ | $RTONF \cdot log(\frac{Nao}{Nai})$ | Equilibrium potential for $Na^{+}$ |
| $E_{mh}$ | – | $RTONF \cdot log(\frac{Nao+0.12 \cdot Ko}{Nai+0.12 \cdot Ki})$ | Equilibrium potential for fast $Na^{+} channel$ |
| $E_{K}$ | $RTONF \cdot log(\frac{Ko}{Ki})$ | $RTONF \cdot log(\frac{Ko}{Ki})$ | Equilibrium potential for $K^{+}$ |
| $E_{Ks}$ | – | $RTONF \cdot log(\frac{Ko+0.12 \cdot Nao}{Ki+0.12 \cdot Nai})$ | Equilibrium potential for slow rectifier $K^{+} channel$ |
| $E_{Ca}$ | $RTONF \cdot log(\frac{Cao}{Cai})$ | $0.5 \cdot RTONF \cdot log(\frac{Cao}{Ca_{sub}})$ | Equilibrium potential for $Ca^{2+}$ |

#### 1.5 Sarcolemmal Ion Currents and Conductances

| | Severi model | Fabbri model | Description |
|---|---|---|---|
| $I_f$ | | | Hyperpolarisation-activated current |
| $g_{f,Na}$ | $0.03\,\mu S$ | $0.00268\,\mu S$ | |
| $g_{f,Na}$ | $0.03\,\mu S$ | $0.00159\,\mu S$ | |
| $Km_f$ | $45\,mM$ | $-$ | |
| $K_{if}$ | $26.26\,mV$ | $26.26\,mV$ | Maximal If activation by cAMP (adjusted based on SLSQP optimisation) |
| $K05_{if}$ | $\frac{17.8741}{600}\,mM$ | $\frac{17.8741}{600}\,mM$ | Half-maximal If activation of cAMP (two step adjustment based on SLSQP optimisation) |
| $n_{if}$ | $9.281$ | $9.281$ | Hill coefficient |
| $V_{if,05}$ | $-52.5\,mV$ | $-$ | Half activation voltage for If current in the basal state |
| $I_{CaL}$ | $P_{CaL} = 0.2\,\frac{nA}{mM}$ | $P_{CaL} = 0.4578\,\frac{nA}{mM}$ | L-type $Ca^{2+}$ current |
| $I_{CaT}$ | $P_{CaT} = 0.02\,\frac{nA}{mM}$ | $P_{CaT} = 0.04132\,\frac{nA}{mM}$ | T-type $Ca^{2+}$ |
| $I_{Kr}$ | $g_{Kr,max} = 0.0021637\,\mu S$ | $g_{Kr} = 0.00424\,\mu S$ | Delayed rectifier $K^+$ current, rapid component |
| $I_{Kur}$ | $-$ | $g_{Kur} = 0.0001539\,\mu S$ | Delayed rectifier $K^+$ current, ultrarapid component |
| $I_{Ks}$ | $g_{Ks,max} = 0.0016576\,\mu S$ | $g_{Ks} = 0.00065\,\mu S$ | Delayed rectifier $K^+$ current, slow component |
| $I_{KACh}$ | $g_{KACh,max} = 0.00864\,\mu S$ | $g_{KACh} = 0.00345\,\mu S$ | ACh-activated $K^+$ current |
| $I_{to}$ | $g_{to,max} = 0.002\,\mu S$ | $g_{to} = 0.0035\,\mu S$ | Transient outward $K^+$ current |
| $I_{Na}$ | $g_{Na,max} = 0.0125\,\mu S$ | $g_{Na} = 0.0223\,\mu S$ | (Fast) $Na^+$ current |
| $I_{NaK}$ | $I_{NaK,max} = 0.063\,nA$ | $I_{NaK} = 0.08105\,nA$ | $Na^+/K^+$ pump current |
| $I_{NaCa}$ | $K_{NaCa} = 4\,nA$ | $K_{NaCa} = 3.343\,nA$ | $Na^+/Ca^{2+}$ exchanger current |
| $\delta_m$ | $1e^{-5}\,mV$ | $1e^{-5}\,mV$ | |

#### 1.6 Modulation of Sarcolemmal Ion Currents by Ions

| | Severi model | Fabbri model | Description |
|---|---|---|---|
| $Km_{fCa}$ | $0.00035\,mM$ | $0.000338\,mM$ | Dissociation constant of $Ca^{2+}$-dependant $I_{CaL}$ inactivation |
| $Km_{Kp}$ | $1.4\,mM$ | $1.4\,mM$ | Half-maximal $Ko$ for $I_{NAK}$ |
| $Km_{Nap}$ | $14.0\,mM$ | $14.0\,mM$ | Half-maximal $Nai$ for $I_{NAK}$ |
| $\alpha_{fCa}$ | $0.01\,\frac{1}{ms}$ | $0.0075\,\frac{1}{s}$ | $Ca^{2+}$ dissociation rate constant for $I_{CaL}$ |

#### 1.7 Na+/Ca2+ Exchanger (NaCa) Function

| | Severi model | Fabbri model | Description |
|---|---|---|---|
| $K1ni$ | $395.3\,mM$ | $395.3\,mM$ | Intracellular $Na^+$ binding to first site on $NaCa$ |
| $K1no$ | $1628.0\,mM$ | $1628.0\,mM$ | Extracellular $Na^+$ binding to first site on $NaCa$ |
| $K2ni$ | $2.289\,mM$ | $2.289\,mM$ | Intracellular $Na^+$ binding to second site on $NaCa$ |
| $K2no$ | $561.4\,mM$ | $561.4\,mM$ | Extracellular $Na^+$ binding to second site on $NaCa$ |
| $K3ni$ | $26.44\,mM$ | $26.44\,mM$ | Intracellular $Na^+$ binding to third site on $NaCa$ |
| $K3no$ | $4.663\,mM$ | $4.663\,mM$ | Extracellular $Na^+$ binding to third site on $NaCa$ |
| $Kci$ | $0.0207\,mM$ | $0.0207\,mM$ | Intracellular $Ca^{2+}$ binding to $NaCa$ transporter |
| $Kcni$ | $26.44\,mM$ | $26.44\,mM$ | Intracellular $Na^+$ and $Ca^{2+}$ simultaneous binding to $NaCa$ |
| $Kco$ | $3.663\,mM$ | $3.663\,mM$ | Extracellular $Ca^{2+}$ binding to $NaCa$ transporter |
| $Qci$ | $0.1369$ | $0.1369$ | Intracellular $Ca^{2+}$ occlusion reaction of $NaCa$ |
| $Qco$ | $0$ | $0$ | Extracellular $Ca^{2+}$ occlusion reaction of $NaCa$ |
| $Qn$ | $0.4315$ | $0.4315$ | $Na^+$ occlusion reactions of $NaCa$ |

#### 1.8 Ca2+ Diffusion

| | Severi model | Fabbri model | Description |
|---|---|---|---|
| $\tau_{dif,Ca}$ | $0.04\,ms$ | $5.469 \cdot 10^{-5}\,s$ | Time constant of $Ca^{2+}$ diffusion from the submembrane to myoplasm |
| $\tau_{tr}$ | $40\,ms$ | $0.04\,s$ | Time constant for $Ca^{2+}$ transfer from the nSR to jSR |

#### 1.9 SR Ca2+ ATPase Function

| | Severi model | Fabbri model | Description |
|---|---|---|---|
| $K_{up}$ | $0.0006\,mM$ | $286\,nM$ | Half-maximal $Cai$ for $Ca^{2+}$ uptake in the nSR |
| $P_{up,basal}$ | $0.012\,\frac{mM}{ms}$ | $5\,\frac{mM}{s}$ | Rate constant for $Ca^{2+}$ uptake by the $Ca^{2+}$ pump in the nSR |
| $slope_{up}$ | – | $50\,nM$ | Slope factor for $Ca^{2+}$ uptake by SERCA pump into the network SR |

#### 1.10 RyR Function

| | Severi model | Fabbri model | Description |
|---|---|---|---|
| $RyR_{min}$ | $0.0127$ | $0.0127$ | (derived from PP1 activity) |
| $RyR_{max}$ | $0.02$ | $0.02$ | |
| $k05_{RyR}$ | $0.682891$ | $0.682891$ | (adjustment based on SLSQP optimisation) |
| $n_{RyR}$ | $9.733$ | $9.733$ | |
| $kiCa$ | $0.5\,\frac{1}{mM\cdot ms}$ | $500\,\frac{1}{mM\cdot s}$ | |
| $kim$ | $0.005\,\frac{1}{ms}$ | $5\,\frac{1}{s}$ | |
| $koCa_{max}$ | $10.0\,\frac{1}{mM^2\cdot ms}$ | $10000.0\,\frac{1}{mM^2\cdot s}$ | (constant in Severi/Fabbri et al. model) |
| $kom$ | $0.06\,\frac{1}{ms}$ | $660\,\frac{1}{s}$ | |
| $ks$ | $250000\,\frac{1}{ms}$ | $1.48\cdot 10^8\,\frac{1}{s}$ | |
| $EC50_{SR}$ | $0.45\,mM$ | $0.45\,mM$ | |
| $HSR$ | $2.5$ | $2.5$ | |
| $MaxSR$ | $15$ | $15$ | |
| $MinSR$ | $1$ | $1$ | |

#### 1.11 $Ca^{2+}$ and $Mg^{2+}$ Buffering

| | Severi model | Fabbri model | Description |
|---|---|---|---|
| $CM_{tot}$ | $0.045\,mM$ | $0.045\,mM$ | Total calmodulin concentration |
| $CQ_{tot}$ | $10.0\,mM$ | $10.0\,mM$ | Toltal calsequestrin concentration |
| $TC_{tot}$ | $0.031\,mM$ | $0.031\,mM$ | Total concentration of the troponin-$Ca^{2+}$ site |
| $TMC_{tot}$ | $0.062\,mM$ | $0.062\,mM$ | Total concentration of the troponin-$Mg^{2+}$ site |
| $kb_{CM}$ | $0.542\,\frac{1}{ms}$ | $542.0\,\frac{1}{s}$ | $Ca^{2+}$ dissociation constant for calmodulin |
| $kb_{CQ}$ | $0.445\,\frac{1}{ms}$ | $445.0\,\frac{1}{s}$ | $Ca^{2+}$ dissociation constant for calsequestrin |
| $kb_{TC}$ | $0.446\,\frac{1}{ms}$ | $446.0\,\frac{1}{s}$ | $Ca^{2+}$ dissociation constant for troponin-$Ca^{2+}$ site |
| $kb_{TMC}$ | $0.0751\,\frac{1}{ms}$ | $7.51\,\frac{1}{s}$ | $Ca^{2+}$ dissociation constant for troponin-$Mg^{2+}$ site |
| $kb_{TMM}$ | $0.751\,\frac{1}{ms}$ | $751.0\,\frac{1}{s}$ | $Mg^{2+}$ dissociation constant for troponin-$Mg^{2+}$ site |
| $kb_{BAPTA}$ | $0.11938\,\frac{1}{ms}$ | – | |
| $kf_{CM}$ | $227.7\,\frac{1}{mM\cdot ms}$ | $1.642\cdot 10^6\,\frac{1}{mM\cdot s}$ | $Ca^{2+}$ association constant for calmodulin |
| $kf_{CQ}$ | $0.534\,\frac{1}{mM\cdot ms}$ | $175.4\,\frac{1}{mM\cdot s}$ | $Ca^{2+}$ association constant for calsequestrin |
| $kf_{TC}$ | $88.8\,\frac{1}{mM\cdot ms}$ | $88800.0\,\frac{1}{mM\cdot s}$ | $Ca^{2+}$ association constant for troponin |
| $kf_{TMC}$ | $227.7\,\frac{1}{mM^2\cdot ms}$ | $227700.0\,\frac{1}{mM\cdot s}$ | $Ca^{2+}$ association constant for the troponin-$Mg^{2+}$ site |
| $kf_{TMM}$ | $2.277\,\frac{1}{mM\cdot ms}$ | $2277.0\,\frac{1}{mM\cdot s}$ | $Mg^{2+}$ association constant for the troponin-$Mg^{2+}$ site |
| $kf_{BAPTA}$ | $940.0\,\frac{1}{mM\cdot ms}$ | – | |
| $T_{Ca,dynamics}$ | $6.928\cdot 10^3\,ms$ | – | |

#### 1.12 AC-cAMP-PKA Signalling

| | Severi model | Fabbri model | Description |
|---|---|---|---|
| $K_{ACI}$ | $0.016\,\frac{1}{min}$ | $0.016\,\frac{1}{min}$ | Non-$Ca^{2+}$ AC activity |
| $K_{AC}$ | $0.0735\,\frac{1}{min}$ | $0.0735\,\frac{1}{min}$ | Non-$Ca^{2+}$ AC activation |
| $K_{Ca}$ | $0.0000896262\,mM$ | $0.000563995\,mM$ | Maximal $Ca^{2+}$ AC activation |
| $K_{ACCa}$ | $0.000024\,mM$ | $0.000024\,mM$ | Half-maximal $Ca^{2+}$ AC activation |
| $K_{fab}$ | - | 4.646336 | Compensation factor introduced due to different $kb_{CM}$ and $kf_{CM} constants$ |
| $k_{PKA}$ | $15\,\frac{mM}{min}$ | $15\,\frac{mM}{min}$ | Maximal PKA activity |
| $k_{PKA,cAMP}$ | $0.474167\,nM$ | $0.474167\,nM$ | Half-maximal PKA activation |
| $n_{PKA}$ | $5$ | $5$ | Hill coefficient |
| $PKA_{tot}$ | $1\,mM$ | $1\,mM$ | Total amount of PKA (adopted from Yaniv et al. model) |
| $PKI_{tot}$ | $0.3\,mM$ | $0.3\,mM$ | Total amount of PKA inhibitor (adopted from Yaniv et al. model) |

#### 1.13 Phosphorylation Parameters

| | Severi model | Fabbri model | Description |
|---|---|---|---|
| $k_{PLBp}$ | $52.25\,\frac{1}{min}$ | $52.25\,\frac{1}{min}$ | Maximal PLB phosphorylation |
| $n_{PLB}$ | $1$ | $1$ | Hill coefficient |
| $k_{PKA,PLB}$ | $1.610336$ | $1.610336$ | Half-maximal PLB phosphorylation (adjusted based on SLSQP optimisation) |
| $PP1$ | $0.00089\,mM$ | $0.00089\,mM$ | PP1 concentration |
| $k_{PP1}$ | $23575.0\,\frac{1}{mM\cdot min}$ | $23575.0\,\frac{1}{mM\cdot min}$ | Maximal PP1 activity |
| $k_{PP1,PLB}$ | $0.06967$ | $0.7457$ | Half-maximal PP1 activation |

#### 1.14 ATP Parameters

| | Severi model | Fabbri model | Description |
|---|---|---|---|
| $ATP_{i,max}$ | $2.533\,mM$ | $2.533\,mM$ | |
| $k_{ATP}$ | $6142$ | $6142$ | |
| $k_{ATP,05}$ | $6724$ | $6724$ | |
| $cAMP_b$ | $0.03333\,mM$ | $0.03333\,mM$ | Basal cAMP concentration |
| $n_{ATP}$ | $3.36$ | $3.36$ | Hill coefficient |
| $k_{ATP,min}$ | $6034$ | $6034$ | |

##### 1.15 Constants

$Iva_{3uM} = 0$
$Cs_{5nM} = 0$
$ACh = 0$
$Iso_{1uM} = 0$
$Iso_{cas} = 0$
$BAPTA_{10nM} = 0$
$CCh_{cas} = 0$
$PI = 3.14159265358979323846264338327950288 4$

#### 1.16 Initial Values

| | Severi model | Fabbri model |
|---|---|---|
| $V_{init}$ | $-52\,mV$ | $-46.1899\,mV$ |
| $Nai_{init}$ | $7.5\,mM$ | $6.084085\,mM$ |
| $y_{init}$ | $0.181334538702451$ | $0.009508$ |
| $m_{init}$ | $0.440131579215766$ | $0.447724$ |
| $h_{init}$ | $1.3676940140066e^{-5}$ | $0.003058$ |
| $dL_{init}$ | $0.0$ | $0.001921$ |
| $fL_{init}$ | $0.497133507285601$ | $0.846702$ |
| $fCa_{init}$ | $0.697998543259722$ | $0.844449$ |
| $dT_{init}$ | $0.0$ | $0.268909$ |
| $fT_{init}$ | $0.0$ | $0.020484$ |
| $R_{Ca,SR,release,init}/R_1$ | $0.912317231017262$ | $0.9308$ |
| $O_{init}$ | $1.7340201253 \cdot 10^{-7}$ | $6.181512 \cdot 10^{-9}$ |
| $I_{init}$ | $7.86181717518 \cdot 10^{-8}$ | $4.595622 \cdot 10^{-10}$ |
| $RI_{init}$ | $0.21114814551282$ | $0.069199$ |
| $fTMM_{init}$ | $0.501049376634$ | $0.653777$ |
| $fCMi_{init}$ | $0.0373817991524254$ | $0.217311$ |
| $fCMs_{init}$ | $0.054381370046$ | $0.158521$ |
| $fTC_{init}$ | $0.0180519400676086$ | $0.017929$ |
| $fTMC_{init}$ | $0.281244308217086$ | $0.259947$ |
| $fCQ_{init}$ | $0.299624275428735$ | $0.138975$ |
| $Cai_{init}$ | $1 \cdot 10^{-5}\,mM$ | $9.15641 \cdot 10^{-6}\,mM$ |
| $Ca_{nsr,init}$ | $1.05386465080816\,mM$ | $0.435148\,mM$ |
| $Ca_{jsr,init}$ | $0.316762674605\,mM$ | $0.409551\,mM$ |
| $Ca_{sub,init}$ | $1 \cdot 10^{-5}\,mM$ | $6.226104 \cdot 10^{-5}\,mM$ |
| $fBAPTA_{init}$ | $0.0$ | $395.3\,mM$ |
| $fBAPTA_{sub,init}$ | $0.0$ | $1628.0\,mM$ |
| $q_{init}$ | $0.506139850982478$ | $0.430836$ |
| $r_{init}$ | $0.0144605370597924$ | $561.4\,mM$ |
| $paS_{init}$ | $0.322999177802891$ | $0.283185$ |
| $paF_{init}$ | $0.0990510403258968$ | $0.011068$ |
| $piy_{init}$ | $0.705410877258545$ | $0.709051$ |
| $n_{init}$ | $0.0$ | $0.1162$ |
| $a_{init}$ | $0.0$ | $0.00277$ |
| $cAMP_{init}$ | $0.032883333\,mM$ | $0.032883333\,mM$ |
| $PLBp_{init}$ | $0.23$ | $0.23$ |
| $r_{to,init}$ | $-$ | $0.014523$ |
| $r_{Kur,init}$ | $-$ | $0.011845$ |
| $s_{Kur,init}$ | $-$ | $0.845304$ |
| $x_{init}$ | $-$ | $0.0685782388661923$ |

### 2 Equations

#### 2.1 Membrane Potential

$\frac{dV}{dt} = \frac{-I_{tot}}{C}$

$I_{tot} = I_f + I_{Kr} + I_{Ks} + I_{to} + I_{NaK} + I_{NaCa} + I_{Na} + I_{CaL} + I_{CaT} + I_{KACh}$

#### 2.2 Ion Currents

$x_\infty$: Steady-state curve for gating variable x
$\tau_x$: Time constant for gating variable x
$\alpha_x$ and $\beta_x$: Opening and closing rates for channel gating

#### 2.3 Hyperpolarisation-activated "funny" Current ($I_f$)

$I_f = I_{f,Na} + I_{f,K}$

$I_{f,Na} = \frac{y^2 \cdot K_o}{K_o + Km_f} \cdot g_{f,Na} \cdot V - E_{Na}$

$I_{f,K} = \frac{y^2 \cdot K_o}{K_o + Km_f} \cdot g_{f,K} \cdot V - E_K$

$Km_f = 45mM$

$Iso_{shift,cas} = \frac{K_{if} \cdot [cAMP]^{n_{if}}}{K_{05,if}^{n_{if}} + [cAMP]^{n_{if}}} - 18.76$

$y_\infty = \frac{1}{1 + e^{\frac{V + 52.5 - Iso_{shift,cas}}{9}}}$

$\tau_y = \frac{0.7}{0.0708 + e^{\frac{-(V+5)}{20.28}} + 10.6 \cdot e^{\frac{V}{18}}}$

$\frac{dy}{dt} = \frac{y_\infty - y}{\tau_y}$

#### 2.4 L-type $Ca^{2+}$ Current ($I_{CaL}$)

$I_{CaL} = I_{siCa} + I_{siK} + I_{siNa} \cdot (1 + Iso_{inc,cas,ICaL})$

$I_{siCa} = \frac{2 \cdot P_{CaL} \cdot V}{RTONF \cdot \left(1 - e^{\frac{-2 \cdot V}{RTONF}}\right)} \cdot \left(Ca_{sub} - Cao \cdot e^{\frac{-2 \cdot V}{RTONF}}\right) \cdot dL \cdot fL \cdot fCa$

$I_{siK} = \frac{0.000365 \cdot P_{CaL} \cdot V}{\left(1 - e^{\frac{-V}{RTONF}}\right)} \cdot \left(K_i - K_o \cdot e^{\frac{-V}{RTONF}}\right) \cdot dL \cdot fL \cdot fCa$

$I_{siNa} = \frac{0.0000185 \cdot P_{CaL} \cdot V}{RTONF \cdot \left(1 - e^{\frac{-V}{RTONF}}\right)} \cdot \left(Nai - Nao \cdot e^{\frac{-V}{RTONF}}\right) \cdot dL \cdot fL \cdot fCa$

$Iso_{inc,cas,ICaL} = \frac{-0.2152 + 0.470657 \cdot [PKA]^{10.0808}}{0.730287^{10.0808} + \cdot [PKA]^{10.0808}}$

$Iso_{shift,cas,ICaL,dLgate} = -(24.4 \cdot 1.128124 \cdot \frac{[PKA]^{9.281}}{0.661450^{9.281} + [PKA]^{9.281}} - 18.76)$

$Iso_{slope,cas} = -1.4762 \cdot [PKA] + 2.1219$

$dL_\infty = \frac{1}{1 + e^{\frac{-(V + 20.3 - Iso_{shift,cas,ICaL,dLgate})}{Iso_{slope,cas} \cdot 4.2}}}$

$\alpha_{dL} = \frac{-0.02839 \cdot (V + 41.8 - Iso_{shift,cas,ICaL,dLgate})}{e^{\frac{-(V+41.8-Iso_{shift,cas,ICaL,dLgate})}{2.5}} - 1} - \frac{0.0849 \cdot (V + 6.8 - Iso_{shift,cas,ICaL,dLgate})}{e^{\frac{-(V+6.8-Iso_{shift,cas,ICaL,dLgate})}{4.8}} - 1}$

$$\beta_{dL} = \frac{0.01143\cdot(V+1.8-Iso_{shift,cas,ICaL,dLgate})}{e^{\frac{V+1.8-Iso_{shift,cas,ICaL,dLgate}}{2.5}}-1}$$

$$\tau_{dL} = \frac{0.001}{\alpha_{dL}+\beta_{dL}}$$

$$\frac{ddL}{dt} = \frac{dL_{\infty}-dL}{\tau_{dL}}$$

$$fL_{\infty} = \frac{1}{1+e^{\frac{V+37.4}{5.3}}}$$

$$\tau_{fL} = 0.001\cdot\left(44.3+230\cdot e^{-\left(\frac{V+36}{10}\right)^2}\right)$$

$$\frac{dfL}{dt} = \frac{fL_{\infty}-fL}{\tau_{fL}}$$

$$fCa_{\infty} = \frac{Km_{fCa}}{Km_{fCa}+Ca_{sub}}$$

$$\tau_{fCa} = \frac{0.001\cdot fCa_{\infty}}{\alpha_{fCa}}$$

$$\frac{dfCa}{dt} = \frac{fCa_{\infty}-fCa}{\tau_{fCa}}$$

#### 2.5 T-type $Ca^{2+}$ Current ($I_{CaT}$)

$$I_{siCa} = \frac{2\cdot P_{CaT}\cdot V}{RTONF\cdot\left(1-e^{\frac{-2\cdot V}{RTONF}}\right)}\cdot\left(Ca_{sub}-Cao\cdot e^{\frac{-2\cdot V}{RTONF}}\right)\cdot dT\cdot fT$$

$$dT_{\infty} = \frac{1}{1+e^{\frac{-(V+38.3)}{5.5}}}$$

$$\tau_{dT} = \frac{0.001}{1.068\cdot e^{\frac{V+38.3}{30}}+1.068\cdot e^{\frac{-(V+38.3)}{30}}}$$

$$\frac{ddT}{dt} = \frac{dT_{\infty}-dT}{\tau_{dT}}$$

$$fT_{\infty} = \frac{1}{1+e^{\frac{V+58.7}{3.8}}}$$

$$\tau_{fT} = \frac{1}{16.67\cdot e^{\frac{-(V+75)}{83.3}}+16.67\cdot e^{\frac{V+75}{15.38}}}$$

$$\frac{dfT}{dt} = \frac{fT_{\infty}-fT}{\tau_{fT}}$$

#### 2.6 Rapidly Activating Delayed Rectifier $K^{+}$ Current ($I_{Kr}$)

$$I_{Kr} = g_{Kr}\cdot(V-E_K)\cdot(0.9\cdot paF+0.1\cdot paS)\cdot piy$$

$$pa_{\infty} = \frac{1}{1+e^{\frac{-(V+14.8)}{8.5}}}$$

$$\tau_{paF} = \frac{1}{30\cdot e^{\frac{V}{10}}+e^{\frac{-V}{12}}}$$

$$\frac{dpaF}{dt} = \frac{pa_{\infty}-paF}{\tau_{paF}}$$

$$\tau_{paS} = \frac{0.84655}{4.2\cdot e^{\frac{V}{17}}+0.15\cdot e^{\frac{-V}{21.6}}}$$

$$\frac{dpaS}{dt} = \frac{pa_{\infty}-paS}{\tau_{paS}}$$

$pi_{\infty} = \frac{1}{1+e^{\frac{V+28.6}{17.1}}}$

$\tau_{pi} = \frac{1}{100 \cdot e^{\frac{-V}{54.645}} + 656 \cdot e^{\frac{V}{106.157}}}$

$\frac{dpi}{dt} = \frac{pi_{\infty} - piy}{\tau_{pi}}$

#### 2.7 Slowly Activating Delayed Rectifier K$^+$ Current ($I_{Ks}$)

$I_{Ks} = (1 + Iso_{inc,cas,IKs}) \cdot g_{Ks} \cdot (V - E_K) \cdot n^2$

$Iso_{inc,cas,IKs} = \frac{-0.2152 + 0.435692 \cdot [PKA]^{10.0808}}{0.704217^{10.0808} + [PKA]^{10.0808}}$

$Iso_{shift,cas,IKs,ngate} = -(24.4 \cdot 1.411375 \cdot \frac{[PKA]^{9.281}}{0.704217^{9.281} + [PKA]^{9.281}} - 18.76)$

$n_{\infty} = \frac{\frac{14}{1+e^{\frac{-(V-40-Iso_{shift,cas,IKs,ngate})}{9}}}}{\frac{14}{1+e^{\frac{-(V-40-Iso_{shift,cas,IKs,ngate})}{9}}} + 1 \cdot e^{\frac{-V}{45}}}$

$\tau_n = \frac{1}{\frac{28}{1+e^{\frac{-(V-40)}{3}}} + e^{\frac{-(V-5)}{25}}}$

$\alpha_n = \frac{\frac{28}{1+e^{\frac{-(V-40-Iso_{shift,cas,IKs,ngate})}{3}}}}{1000}$

$\beta_n = \frac{e^{\frac{-(V-5-Iso_{shift,cas,IKs,ngate})}{25}}}{1000}$

$\frac{dn}{dt} = \frac{n_{\infty} - n}{\tau_n}$

#### 2.8 ACh-Activated K$^+$ Current ($I_{KACh}$)

$$I_{KACh} = \begin{cases} g_{KACh} \cdot (V - E_K) \cdot \left(1 + e^{\frac{V+20}{20}}\right) \cdot a, & if\ ACh > 0 \\ 0, & otherwise \end{cases}$$

$a_{\infty} = \frac{\alpha_a}{\alpha_a + \beta_a}$

$\alpha_a = \frac{3.5988 - 0.0256}{1 + \frac{0.0000012155}{ACh^{1.6951}}} + 0.0256$

$\beta_a = 10 \cdot e^{0.0133 \cdot (V+40)}$

$\tau_a = \frac{1}{\alpha_a + \beta_a}$

$\frac{da}{dt} = \frac{a_{infty} - a}{\tau_{\alpha}}$

#### 2.9 Transient Outward K$^+$ Current ($I_{to}$)

$I_{to} = g_{to} \cdot (V - E_K) \cdot q \cdot r$

$q_{\infty} = \frac{1}{1+e^{\frac{V+49}{13}}}$

$\tau_q = 0.001 \cdot 0.6 \cdot \left(\frac{65.17}{0.57 \cdot e^{-0.08 \cdot (V+44)} + 0.065 \cdot e^{0.1 \cdot (V+45.93)}} + 10.1\right)$

$\frac{dq}{dt} = \frac{q_\infty - q}{\tau_q}$

$r_\infty = \frac{1}{1+e^{\frac{-(V-19.3)}{15}}}$

$\tau_r = 0.001 \cdot 0.66 \cdot 1.4 \cdot \left(\frac{15.59}{1.037 \cdot e^{0.09\cdot(V+30.61)} + 0.369 \cdot e^{-0.12\cdot(V+23.84)}} + 2.98\right)$

$\frac{dr}{dt} = \frac{r_\infty - r}{\tau_r}$

#### 2.10 Na$^+$ Current ($I_{Na}$)

$I_{Na} = g_{Na} \cdot (V - E_{mh}) \cdot m^3 \cdot h$

$E_{mh} = RTONF \cdot log\left(\frac{Nao+0.12\cdot Ko}{Nai+0.12\cdot Ki}\right)$

$E0_m = V + 41$

$\delta_m = 1 \cdot 10^{-5} mV$

$\frac{dm}{dt} = \alpha_m \cdot (1-m) - \beta_m \cdot m$

$$\alpha_m = \begin{cases} 2000, & if\,|E0_m| < \delta_m \\ \frac{200\cdot E0_m}{1-e^{-0.1\cdot E0_m}}, & otherwise \end{cases}$$

$\beta_m = 8000 \cdot e^{-0.056\cdot(V+66)}$

$\frac{dh}{dt} = \alpha_h \cdot (1-h) - \beta_h \cdot h$

$\alpha_h = 20 \cdot e^{-0.125\cdot(V+75)}$

$\beta_h = \frac{2000}{320\cdot e^{-0.1\cdot(V+75)}+1}$

#### 2.11 Na$^+$-K$^+$ Pump Current ($I_{NaK}$)

$I_{NaK} = (1 + Iso_{inc,cas,INaK}) \cdot I_{NaK,max} \cdot \left(1+\left(\frac{Km_{Kp}}{Ko}\right)^{1.2}\right)^{-1} \cdot \left(1+\left(\frac{Km_{Nap}}{Nai}\right)^{1.3}\right)^{-1} \cdot \left(1+e^{\frac{-(V-E_{Na}+100)}{20}}\right)^{-1}$

$Iso_{inc,cas,INaK} = -0.2152 + 0.435692 \cdot \frac{[PKA]^{10.0808}}{0.719701^{10.0808}+[PKA]^{10.0808}}$

#### 2.12 Na$^+$-Ca$^{2+}$ Exchanger Current ($I_{NaCa}$)

$I_{NaCa} = \frac{K_{NaCa}\cdot(x2\cdot k21 - x1\cdot k12)}{x1+x2+x3+x4}$

$x1 = k41 \cdot k34 \cdot (k23 + k21) + k21 \cdot k32 \cdot (k43 + k41)$

$x2 = k32 \cdot k43 \cdot (k14 + k12) + k41 \cdot k12 \cdot (k34 + k32)$

$x3 = k14 \cdot k43 \cdot (k23 + k21) + k12 \cdot k23 \cdot (k43 + k41)$

$x4 = k23 \cdot k34 \cdot (k14 + k12) + k14 \cdot k21 \cdot (k34 + k32)$

$k43 = \frac{Nai}{K3ni+Nai}$

$k12 = \frac{\frac{Ca_{sub}}{Kci} \cdot e^{\frac{-Qci \cdot V}{RTONF}}}{di}$

$k14 = \frac{\frac{\frac{Nai}{K1ni} \cdot Nai}{K2ni} \cdot (1+\frac{Nai}{K3ni}) e^{\frac{Qn \cdot V}{2 \cdot RTONF}}}{di}$

$k41 = e^{\frac{-Qn \cdot V}{2 \cdot RTONF}}$

$di = 1 + \frac{Ca_{sub}}{Kci} \cdot \left(1 + e^{\frac{-Qci \cdot V}{RTONF}} + \frac{Nai}{Kcni}\right) + \frac{Nai}{K1ni} \cdot \left(1 + \frac{Nai}{K2ni} \cdot \left(1 + \frac{Nai}{K3ni}\right)\right)$

$k34 = \frac{Nao}{K3no+Nao}$

$k21 = \frac{\frac{Ca_o}{Kco} \cdot e^{\frac{Qco \cdot V}{RTONF}}}{do}$

$k23 = \frac{\frac{\frac{Nao}{K1no} \cdot Nao}{K2no} \cdot (1+\frac{Nao}{K3no}) e^{\frac{-Qn \cdot V}{2 \cdot RTONF}}}{do}$

$k23 = e^{\frac{Qn \cdot V}{2 \cdot RTONF}}$

$do = 1 + \frac{Ca_o}{Kco} \cdot \left(1 + e^{\frac{Qco \cdot V}{RTONF}}\right) + \frac{Nao}{K1no} \cdot \left(1 + \frac{Nao}{K2no} \cdot \left(1 + \frac{Nao}{K3no}\right)\right)$

#### 2.13 $Ca^{2+}$ Release Flux ($J_{rel}$) from SR via RyRs

$J_{rel} = ks \cdot O \cdot (Ca_{jsr} - Ca_{sub})$

$kCaSR = MaxSR - \frac{MaxSR-MinSR}{1+\left(\frac{EC50_{SR}}{Ca_{jsr}}\right)^{HSR}}$

$koCa = koCa_{max} \cdot (RyR_{min} - RyR_{max} \cdot \frac{[PKA]^{n_{RyR}}}{k_{05,RyR}^{n_{RyR}}+[PKA]^{n_{RyR}}} - 1)$

$koSRCa = \frac{koCa}{kCaSR}$

$kiSRCa = kiCa \cdot kCaSR$

$\frac{R}{dt} = kim \cdot RI - kiSRCa \cdot Ca_{sub} \cdot R - (koSRCa \cdot Ca_{sub}^2 \cdot R - kom \cdot O)$

$\frac{O}{dt} = koSRCa \cdot Ca_{sub}^2 \cdot R - kom \cdot O - (kiSRCa \cdot Ca_{sub} \cdot O - kim \cdot I)$

$\frac{I}{dt} = kiSRCa \cdot Ca_{sub} \cdot O - kim \cdot I - (kom \cdot I - koSRCa \cdot Ca_{sub}^2 \cdot RI)$

$\frac{RI}{dt} = kom \cdot I - koSRCa \cdot Ca_{sub}^2 \cdot RI - (kim \cdot RI - kiSRCa \cdot Ca_{sub} \cdot R)$

#### 2.14 Intracellular $Ca^{2+}$ Fluxes

$J_{diff}$: $Ca^{2+}$ diffusion flux from submembrane space to myoplasm

$J_{tr}$: $Ca^{2+}$ transfer flux from network to junctional SR

$J_{up}$: $Ca^{2+}$ uptake by SR

$J_{diff} = \frac{Ca_{sub} - Cai}{\tau_{dif,Ca}}$

$J_{tr} = \frac{Ca_{nsr} - Ca_{jsr}}{\tau_{tr}}$

$$F_{PLBp} = \begin{cases} 3.3931 \cdot \frac{[PLBp]^{4.0695}}{0.2805^{4.0695} + [PLBp]^{4.0695}}, & if\ [PLBp] > 0.23 \\ 1.698 \cdot \frac{[PLBp]^{13.5842}}{0.2240^{13.5842} + [PLBp]^{13.5842}}, & otherwise \end{cases}$$

$J_{up} = \frac{0.9 \cdot P_{up} \cdot F_{PLBp}}{1 + \frac{K_{up}}{Cai}}$

#### 2.15 Ca$^{2+}$ Buffering

$f_{CMi}$: Fractional occupancy of calmodulin by Ca$^{2+}$ in myoplasm

$f_{CMi}$: Fractional occupancy of calmodulin by Ca$^{2+}$ in myoplasm

$f_{CMs}$: Fractional occupancy of calmodulin by Ca$^{2+}$ in subspace

$f_{CQ}$: Fractional occupancy of calsequestrin by Ca$^{2+}$

$f_{TC}$: Fractional occupancy of the troponin-Ca$^{2+}$ site by Ca$^{2+}$

$f_{TMC}$: Fractional occupancy of the troponin-Mg$^{2+}$ site by Ca$^{2+}$

$f_{TMM}$: Fractional occupancy of the troponin-Mg$^{2+}$ site by Mg$^{2+}$

$\frac{dfCMi}{dt} = \delta_{fCMi}$

$\delta_{fCMi} = kf_{CM} \cdot Cai \cdot (1 - fCMi) - kb_{CM} \cdot fCMi$

$\frac{dfCMs}{dt} = \delta_{fCMs}$

$\delta_{fCMs} = kf_{CM} \cdot Ca_{sub} \cdot (1 - fCMs) - kb_{CM} \cdot fCMs$

$\frac{dfCQ}{dt} = \delta_{fCQ}$

$\delta_{fCQ} = kf_{CQ} \cdot Ca_{jsr} \cdot (1 - fCQ) - kb_{CQ} \cdot fCQ$

$\frac{dfTC}{dt} = \delta_{fTC}$

$\delta_{fTC} = kf_{TC} \cdot Cai \cdot (1 - fTC) - kb_{TC} \cdot fTC$

$\frac{dfTMC}{dt} = \delta_{fTMC}$

$\delta_{fTMC} = kf_{TMC} \cdot Cai \cdot (1 - (fTMC + fTMM)) - kb_{TMC} \cdot fTMC$

$\frac{dfTMM}{dt} = \delta_{fTMM}$

$\delta_{fTMM} = kf_{TMM} \cdot Mgi \cdot (1 - (fTMC + fTMM)) - kb_{TMM} \cdot fTMM$

#### 2.16 Daynamics of Ca$^{2+}$ Concentration in Cell Compartments

$$\frac{dCai}{dt} = \left(\frac{J_{diff}\cdot V_{sub} - J_{up}\cdot V_{nsr}}{V_i} - (CM_{tot}\cdot\delta_{CMi} + TC_{tot}\cdot\delta_{fTC} + TMC_{tot}\cdot\delta_{fTMC})\right) - \frac{dfBAPTA}{dt}$$

$$\frac{dCa_{sub}}{dt} = \left(\frac{J_{rel}\cdot V_{jsr}}{V_{sub}} - \left(\frac{I_{siCa} + I_{CaT} - 2\cdot I_{NaCa}}{2\cdot F\cdot V_{sub}} + J_{Ca,dif}\cdot CM_{tot}\cdot\delta_{CMs}\right)\right) - \frac{dfBAPTA_{sub}}{dt}$$

$$\frac{dCa_{nsr}}{dt} = J_{up} - \frac{J_{tr}\cdot V_{jsr}}{V_{nsr}}$$

$$\frac{dCa_{jsr}}{dt} = J_{tr} - (J_{rel} + CQ_{tot}\cdot\delta_{fCQ})$$

#### 2.17 Daynamics of Intracellular Na$^{+}$ Concentration

$$\frac{Nai}{dt} = -\frac{I_{Na} + I_{fNa} + I_{siNa} + 3\cdot I_{NaK} + 3\cdot I_{NaCa}}{(V_i + V_{sub})\cdot F}$$

#### 2.18 AC-cAMP-PKA Signalling

$$\frac{d[cAMP]}{dt} = \frac{(k_{ISO} - k_{CCh})\cdot[ATP]_i + k1\cdot[ATP]_i - k2\cdot[cAMP] - k3\cdot[cAMP]}{60000}$$

The factor 60000 converts minutes into milliseconds.

$$k_{ISO} = 0.007 + 0.1181\cdot\frac{[Iso_{cas}]^{0.8664}}{48.1212^{0.8664} + [Iso_{cas}]^{0.8664}}$$

$$k_{CCh} = 0.0146\cdot\frac{[CCh_{cas}]^{1.4402}}{51.7331^{1.4402} + [CCh_{cas}]^{1.4402}}$$

$$k1 = K_{ACI} + \frac{K_{AC}}{1 + e^{\left(\frac{K_{Ca} - \frac{kb_{CM}\cdot fCMi}{kf_{CM}\cdot(1 - fCMi)}}{K_{ACCa}}\right)}}$$

$$k2 = 1.1\cdot 237.9851\cdot\frac{([cAMP]\cdot 600)^{5.101}}{20.1077^{6.101} + ([cAMP]\cdot 600)^{6.101}}$$

The factor 600 converts mM to pM/mg protein.

$$k3 = k_{PKA}\cdot\frac{[cAMP]^{n_{PKA}-1}}{k_{PKA,cAMP}^{n_{PKA}} + [cAMP]^{n_{PKA}}}$$

#### 2.19 PLB Activity

$$\frac{d[PLBp]}{dt} = \frac{k4 - k5}{60000}$$

The factor 60000 converts minutes into milliseconds.

$$k4 = \frac{k_{PLBp}\cdot[PKA]^{n_{PLB}}}{k_{PKA,PLB}^{n_{PLB}} + [PKA]^{n_{PLB}}}$$

$$k5 = \frac{k_{PP1}\cdot PP1\cdot PLBp}{k_{PP1,PLB} + [PLBp]}$$

#### 2.20 PKA Activity

$$[PKA] = \begin{cases} -0.9483029\cdot e^{-[cAMP]\cdot 600\cdot 0.06561479} + 0.97781646, & if\ [cAMP]\cdot 600 < 25.87 \\ \\ -0.45260509\cdot e^{-[cAMP]\cdot 600\cdot 0.03395094} + 0.99221714, & otherwise \end{cases}$$

The factor 600 converts mM to pM/mg protein.

#### 2.21 ATP

$$[APT]_i = \frac{ATP_{i.max}\left(\cdot\frac{k_{ATP}\cdot\left(\frac{[cAMP]\cdot 100}{cAMP_b}\right)^{n_{ATP}}}{k_{ATP,05}+\left(\frac{[cAMP]\cdot 100}{cAMP_b}\right)^{n_{ATP}}}-K_{ATP,min}\right)}{100}$$
