## Supplementary Material - Code for "Sympathetic Stimulation Can Compensate for Hypocalcaemia-Induced Bradycardia in Human and Rabbit Sinoatrial Node Cells": METADATA.yml

```
# According to the DataCite 4.0 scheme: http://doi.org/10.5438/0012
# keyword-only entries need to be supplied by the user
title: # add the title of your experiment
keywords: # more can be added
- cardiac electrophysiology
- openCARP experiment
- in-silico medicine
- computational cardiology
- sinoatrial node cell models
descriptions:
- description: This experiment offers the possibility to simulate the extended sinoatrial node cell models of Severi et al. (2012) and Fabbri et al. (2017) with several Cao and ISO concentrations. The extension of the AC-cAMP-PKA signalling cascade is based on the work of Behar et al. (2016).
  description_type: Abstract
subjects: # more can be added
- subject: Computer simulation
  value_uri: http://id.loc.gov/authorities/subjects/sh85029533
  scheme_uri: http://id.loc.gov/authorities/subjects
- subject: Biomedical engineering
  value_uri: http://id.loc.gov/authorities/subjects/sh85014237
  scheme_uri: http://id.loc.gov/authorities/subjects
- subject: Cardiology
  value_uri: http://id.loc.gov/authorities/subjects/sh85020214
  scheme_uri: http://id.loc.gov/authorities/subjects
radar_subjects:
- LifeScience
- ComputerScience
- Medicine
creators:
- name:
  given_name: Moritz
  family_name: Linder
  orcid: 0009-0000-1228-159X
  affiliations:
  - name: Institute of Biomedical Engineering, Karlsruhe Institute of Technology (KIT), Karlsruhe, Germany
    ror: https://ror.org/04t3en479
publisher: openCARP
publication_year: '2024' # to be auto-populated during bundle creation
date: '2024-08-22' # to be auto-populated during bundle creation YYYY-MM-DD
date_type: collected
resource: Simulation code
resource_type: Workflow
related_identifiers: # add IsNewVersionOf IsDocumentedBy IsSupplementTo as desired
- relation_type: IsCompiledBy
  related_identifier: 10.35097/562
  related_identifier_type: DOI
- relation_type: References
  related_identifier: 10.1016/j.cmpb.2021.106223
  related_identifier_type: DOI
rights: Apache License 2.0
rights_url: https://git.opencarp.org/openCARP/experiment-template/-/blob/master/LICENSE
rights_holder: Moritz Linder # to be autopopulated during bundle creation
funding_references:
- name: European High-Performance Computing Joint Undertaking EuroHPC (JU)
  ror: https://ror.org/00k4n6c32
  award_number: 955495
  award_uri: https://eurohpc-ju.europa.eu/ongoing-projects#ecl-inpage-259
  award_title: Numerical modeling of cardiac electrophysiology at the cellular scale

# Information about software (automatically collected at bundle creation)

bundle_creator_name: Moritz Linder
bundle_creator_email:
bundle_creator_orcid: 0009-0000-1228-159X
python_version: 3.12.3
opencarp_git_hash: v16.0-121-gbd8e10ed
experiment_name: compensatory_effect
experiment_command_line: 'run.py '
```
