## Supplementary Material - Code for "Sympathetic Stimulation Can Compensate for Hypocalcaemia-Induced Bradycardia in Human and Rabbit Sinoatrial Node Cells": run.py

```
#!/usr/bin/env python
import os
from carputils import tools
from random import randint

#### ---------------------------------------------------------------- ####
# This script offers the possibility to simulate the extended sinoatrial 
# node cell models of Severi et al. (2012) and Fabbri et al. (2017) with 
# several [Ca2+]o and [ISO] concentrations. The extension of the AC-cAMP-PKA 
# signalling cascade is based on the work of Behar et al. (2016). To 
# simulate different [Ca2+]o and [ISO] concentrations change the lists in 
# the main function of the script and execute it with the 
# --compensatory-effect flag set to True. To simulate the original models
# without the extension, set [ISO] to -1.
#### ---------------------------------------------------------------- ####

def parser():
    models = ['Severi', 'Fabbri']
    parser = tools.standard_parser()
    group = parser.add_argument_group('Experiment specific options.')
    group.add_argument('--imp', default = models[0], choices = models,
                       help = 'baseline model')
    group.add_argument('--duration', default=1000000.0, type = float,
                       help = 'duration of the experiment [ms] (default=5000.0)')
    group.add_argument('--dt', default=0.01,
                       help = 'time step [ms] (default=0.01)')
    group.add_argument('--dt-out', default = 1.0,
                       help = 'temporal output granularity [ms] (defult=1.0)')
    group.add_argument('--start-out', default=900000.0,
                       help= 'start time of output [ms]  (default=0.0)')
    group.add_argument('--output', default='Trace_0.dat',
                       help = 'output filename')
    group.add_argument('--imp-par', default='',
                       help = 'prams to modify IMP (default='')')
    group.add_argument('--stim-curr', default=0.0,
                       help='stimulation current [pA/pF]=[uA/cm^2] (default=0.0)')
    group.add_argument('--save-ini-time', default=0.0,
                       help='time at which to save single cell state [ms] (default=0.0)')
    group.add_argument('--save-ini-file', default='singlecell.sv',
                       help="text file in which to save state of a single cell (default='singlecell.sv')")
    group.add_argument('--read-ini-file', default='singlecell.sv',
                       help="text file from which to read state of a single cell (default='singlecell.sv')")
    group.add_argument('--compensatory-effect', default=False,
                       help='set to True to simulate several Cao and ISO concentrations defined in main function')
    return parser


# define job ID
def jobID(args):
    """
    Generate name of top level output directory.
    """
    tpl = '{}_{}ms_dt_{}ms'
    return tpl.format(args.imp, args.duration, args.dt)


@tools.carpexample(parser, jobID, clean_pattern='exp*')
def run(args, job):

    if not os.path.exists(f'{job.ID}'):
        os.makedirs(f'{job.ID}')

    # basic simulation settings
    cmd = ['--imp', args.imp]
    cmd += ['--duration', args.duration]
    cmd += ['--dt', args.dt]
    cmd += ['--dt-out', args.dt_out]
    cmd += ['--start-out', args.start_out]
    cmd += ['--stim-curr', args.stim_curr]
    cmd += ['--imp-par', args.imp_par]

    # read or store init files (compensatory experiments)
    # cmd += ['--save-ini-time', args.save_ini_time]
    # cmd += ['--save-ini-file', args.save_ini_file]
    # cmd += ['--read-ini-file', args.read_ini_file]

    # output settings (Trace_$(trace_no).dat to avoid collisions)
    trace_no = randint(0, 1000000)
    cmd += ['--trace-no', trace_no]

    # start the cellular simulation
    title = 'Simulating cellular model {} for {} ms timestep {} ms'
    job.bench(cmd, msg=title.format(args.imp, args.duration, args.dt, args.dt_out, args.imp_par))
    job.mv(os.path.join(f'Trace_{trace_no}.dat'), os.path.join(job.ID, f'{args.imp}_{args.duration}_dt{args.dt}ms_{args.imp_par}'))


if __name__ == '__main__':

    # check for --compensatory effect
    args = parser().parse_args()

    if args.compensatory_effect:
    
        # define Cao and ISO concentrations
        # extended Severi model: Cao [1.3, 1.8]; ISO [0.0, 1000.0]
        # extended Fabbri model: Cao [1.2, 1.8]; ISO [0.0, 1000.0]
        Cao_list = [1.4, 1.6, 1.8]    # change for different [Ca2+]o
        ISO_list = [0.0, 10.0, 100.0, 1000.0]    # change for different [ISO]

        for Cao in Cao_list:
            for ISO in ISO_list:
                run(['--imp', f'{args.imp}', '--imp-par', f'Cao={Cao},Iso_cas={ISO}', '--overwrite-behaviour', 'append'])

    else:
        run()
```
